# Reaction-aware hypergraph learning of chemical representations with LARK

**DOI:** 10.64898/2026.09.15.751932

**Authors:** Jianbo Qiao, Yuhang Liu, Junru Jin, Ding Wang, Quan Zou, Ran Su, Leyi Wei

**Affiliations:** School of Software, Shandong University, Jinan, China; Faculty of Applied Sciences, Macao Polytechnic University, Macao, 999078, China; New Laboratory of Pattern Recognition (NLPR), State Key Laboratory of Multimodal Artificial Intelligence Systems (MAIS), Institute of Automation, Chinese Academy of Sciences; School of Artificial Intelligence, University of Chinese Academy of Sciences; Institute of Fundamental and Frontier Sciences, University of Electronic Science and Technology of China, Chengdu, 610054, China; College of Intelligence and Computing, Tianjin University, Tianjin, 300350, China; Engineering Research Centre of Applied Technology on Machine Translation and Artificial Intelligence, Macao Polytechnic University, Macao SAR, China

## Abstract

Molecular pretraining offers a route to learning transferable chemical representations from unlabelled data. However, existing approaches pretrained on isolated molecular structures struggle to generalize to reaction-specific tasks because their pretraining objectives provide limited information about chemical transformations. A key challenge is to integrate this reaction information into chemical representations. Here we introduce LARK, a unified framework for learning atom–molecule–reaction knowledge from chemical transformations. Using approximately 0.6 million reactions encompassing 0.88 million unique molecular components, LARK combines bidirectional bond–electron reconstruction with structural and geometric supervision through a role-aware hierarchical hypergraph Transformer that links local atom representations to reaction context. The learned representations transfer to both reaction prediction and single-molecule property prediction, with strong performance across 42 tasks covering reaction outcomes, physicochemical properties, toxicity, metabolism and pharmacokinetics. Reaction-centre analyses show that LARK identifies where bonds break and form, while targeted interventions reveal that the learned atom representations contribute to reconstructing these changes. These findings indicate that LARK learns chemically informative representations of substructures involved in molecular transformations. Together, the results establish chemical transformations as a source of structured supervision for representation learning across molecular and reaction scales.

## Introduction

Chemical reactions express the dynamic nature of chemistry through changes in molecular structure, bonding and electron organization. Learning representations that connect molecular structure with chemical behaviour is a central objective of molecular pretraining. By constructing supervision from molecular topology, geometry and associated modalities, pretraining can draw on large collections of unlabelled molecules to learn representations that transfer to downstream property prediction[1–3]. Although these objectives capture structural and physico-chemical regularities, pretraining on isolated molecules provides limited direct supervision about how molecular structures change during reactions. Transfer to reaction-specific tasks therefore requires representations that connect local transformation patterns with the roles and interactions of multiple reaction components.

Reaction-informed learning has already begun to exploit this opportunity. Synthesis knowledge graphs connect molecular representations through reaction relationships, and chemically inspired pretraining links reaction representation learning to conditional molecule generation[4, 5]. Explicit molecular-edit languages make structural transformations accessible to language models, while unified reaction frameworks support both performance prediction and synthesis planning[6, 7]. These advances establish chemical transformations as a useful learning resource. A remaining challenge is to integrate directional local changes with multicomponent reaction context while retaining molecular information that supports property prediction. The changed atoms constitute only a small part of a multicomponent reaction, and substrates, reagents, catalysts and solvents contribute different contextual information. A transferable representation must therefore connect local transformation signals with molecular structure and component roles, while supporting both molecular and reaction inputs.

The bond and valence-electron reorganization central to reaction mechanisms motivates a chemically explicit form of supervision. Electron redistribution has been used to constrain reaction generation and predict mechanistic transformations[8]. For reaction-pair pretraining, bond–electron matrices jointly encode bonding and local electron assignment, and their differences describe the net reorganization between reactants and products. Learning these changes from either side of a reaction requires associating the available molecular structure and reaction context with a directional transformation. This motivates our hypothesis that local change supervision, combined with structural and geometric objectives, can organize atom states around chemically transformed sites while retaining molecular information for transfer. Local chemical changes occur within multicomponent reactions, where the information relevant to a transformation is distributed across molecular structures and component roles. A role-aware reaction hypergraph provides a way to integrate these components within a shared reaction context, while hierarchical feedback connects this context to the atom states used for reconstruction.

Here we introduce LARK, a unified framework for learning atom–molecule–reaction knowledge from chemical transformations. LARK combines bidirectional bond–electron reconstruction with a role-aware hierarchical reaction hypergraph Transformer. A molecular graph encoder first generates atom representations and a summary token for each molecular component. The reaction hypergraph Transformer combines these molecular tokens with component-role embeddings and a reaction token, using self-attention to exchange information across the reaction. Gated feedback then returns the resulting contextual information to atom representations, linking local bond–electron reconstruction with the multicomponent reaction environment. Together with structural and geometric supervision, this architecture supports representation learning at both molecular and reaction scales. We first test whether the resulting representations support diverse reaction predictions and transfer to single-molecule property tasks, including physicochemical properties, bioactivity and absorption, distribution, metabolism, excretion and toxicity[9, 10]. Matched ablations then assess which components support transfer, and reaction-centre localization with targeted interventions tests whether the model uses atom states associated with chemical changes. Together, these evaluations examine chemical transformations as structured supervision for representations useful at both molecular and reaction scales.

## Results

### LARK links local chemical changes with molecular and reaction context

Inspired by the bond and valence-electron reorganization central to reaction mechanisms, LARK uses chemical changes to supervise representation learning (Fig. 1). Bond–electron matrices describe bonding and local electron assignment, and their difference, ΔBE = BE_P_ − BE_R_, specifies the net change between reactants and products. This target connects molecular structure to the atoms and bonds involved in a transformation. Mapped core components define the reconstruction target, while entirely unmapped components provide reaction context (Fig. 1a).

**Figure 1.**
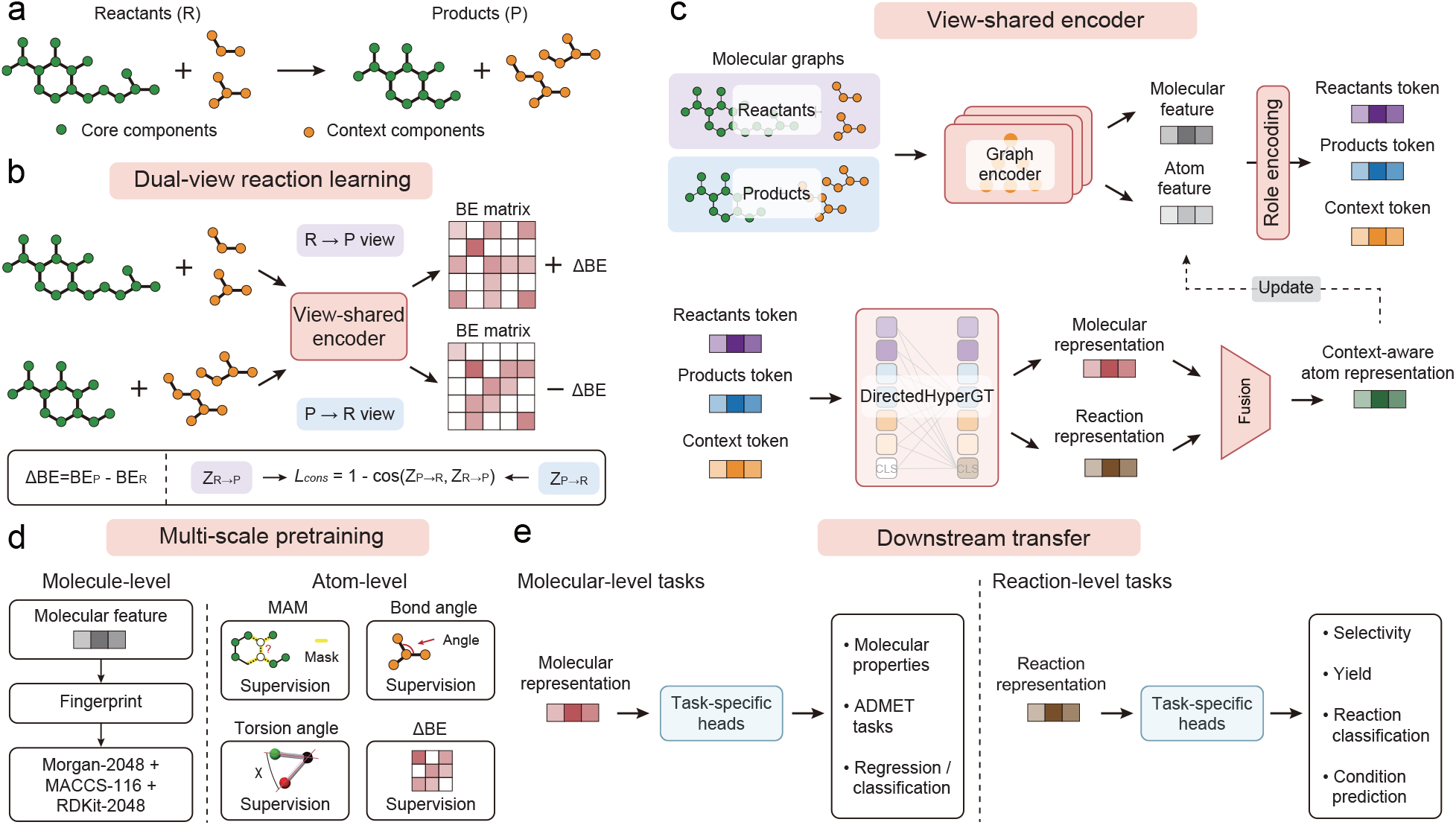
Dual-view bond–electron pretraining and hierarchical transfer in LARK. **a**, A mapped reaction is decomposed into atom-mapped core components and unmapped contextual components on the reactant and product sides. **b**, Complementary reactant-to-product and product-to-reactant views share an encoder, each receiving its source-side mapped core and generic context. The former predicts +ΔBE, whereas the latter predicts *−*ΔBE, where ΔBE = BE_P_ *−* BE_R_. A cosine consistency loss aligns the two view representations. **c**, The view-shared encoder is applied separately to each directional input. A molecular graph encoder produces atom and molecular features. A role-aware reaction hypergraph Transformer (DirectedHyperGT) uses component-role encodings and full self-attention to jointly update molecular tokens and a reaction token. Gated fusion returns reaction-aware molecular context to the atom states. **d**, Five pretraining task types supervise molecular fingerprints, masked atoms, bond angles, torsion angles and directional ΔBE reconstruction. **e**, Molecular and reaction rep-resentations are transferred to molecular-property, ADMET, regression, classification, selectivity, yield, reaction-classification and condition-prediction tasks. The figure schematically defines the architecture and training workflow.

Dual-view reaction learning associates each side of a reaction with its directional change (Fig. 1b). The reactant-to-product (R→P) view receives the mapped reactant core and predicts +ΔBE; the product-to-reactant (P→R) view receives the mapped product core and predicts −ΔBE. Both views receive generic unmapped context and share encoder weights. A cosine consistency objective aligns projected, pooled mapped-core representations, linking the two views while preserving their direction-specific reconstruction targets.

The hierarchy connects these local transformation targets with the molecular components surrounding them (Fig. 1c). A graph encoder first produces atom states and a summary token for each molecule. A role-aware reaction hypergraph Transformer, termed DirectedHyperGT, combines component-role embeddings with full self-attention over molecular tokens and a reaction token. Gated feedback then returns reaction-aware molecular information to local atom states. This arrangement allows reconstruction to draw on both local structure and multicomponent context, while retaining outputs at molecular and reaction scales.

Structural and geometric objectives provide additional supervision for information beyond the changed sites (Fig. 1d). Fingerprint prediction supervises molecular tokens, while masked-atom, bond-angle and torsion-angle prediction accompany directional ΔBE reconstruction at atom level. The resulting framework couples learning about molecular structure with learning about chemical change. Task-specific heads use its molecular or reaction representations for property, selectivity, yield, reaction-class and condition prediction (Fig. 1e).

### LARK supports reaction prediction and few-shot transfer across chemical systems

To test whether representations learned from chemical transformations support diverse reaction predictions, we first evaluated LARK across activation free energies, reactive-site preferences, selectivity and yield (Fig. 2a,b and Supplementary Table S1). On Li/Hong radical C–H functionalization, LARK achieved an *R*^2^ of 0.9828 for absolute activation free energy, 4.7% higher than the reported PhysOrg-RF result for the same prediction target[11]. Transfer also extended to stereoselectivity: relative to Chemma, *R*^2^ was 0.7% higher on Zahrt and 2.4% lower on Long/Ding enantiomeric excess[12]. Prediction plots and residual diagnostics characterize performance across the observed energy and selectivity ranges (Fig. 2c,d). These comparisons support transfer to both activation-energy and stereoselectivity prediction.

**Figure 2.**
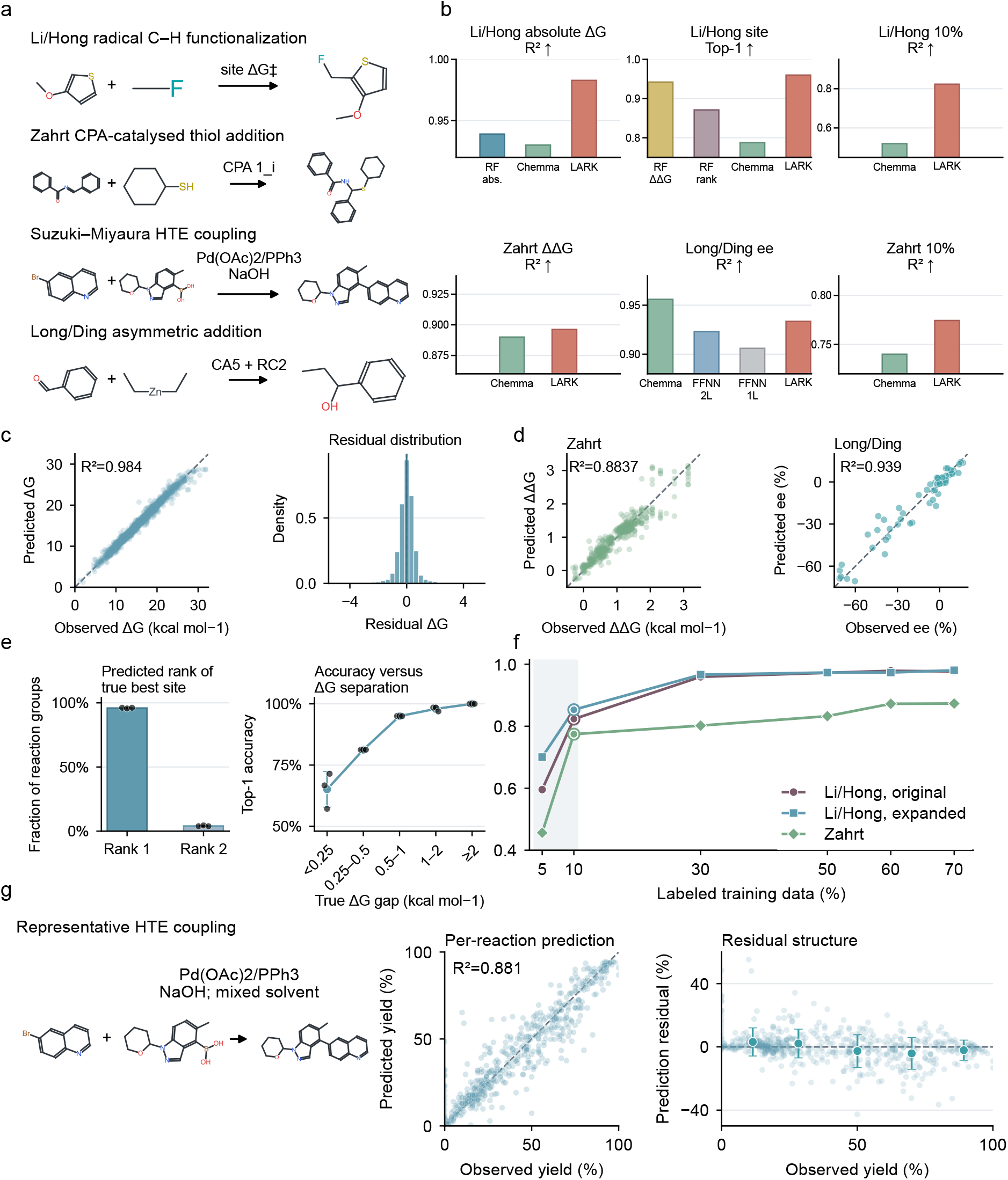
Reaction outcome prediction across chemical systems with limited labelled data. **a**, Representative Li/Hong, Zahrt, Suzuki–Miyaura and Long/Ding reaction systems. **b**, Task-wise comparison with published reference values. The Li/Hong energy comparison includes only absolute Δ*G* prediction; related ΔΔ*G* results are listed separately in Supplementary Table S1. LARK bars are means. Approximate published values retain the approximation symbol. **c**, Observed and predicted Li/Hong absolute Δ*G* with residual distribution (*n* = 1,835 fixed-Test candidates). **d**, Observed and predicted Zahrt ΔΔ*G* (*n* = 475) and Long/Ding ee (*n* = 65, three-fold out-of-fold prediction). **e**, Rank of the true best Li/Hong site and Top-1 accuracy stratified by the true Δ*G* gap. Points are downstream repeats and bars/lines are mean sample *±* s.d. across three seeds; each seed contains 340 reaction groups. **f**, Learning curves for Zahrt and two Li/Hong settings; points are single aggregate Test estimates. **g**, Representative Suzuki HTE reaction, observed–predicted Test yields and residuals grouped by observed yield (*n* = 576); large points and error bars denote within-bin mean sample *±* s.d. Residual is predicted minus observed.

For competing reaction sites, the practical question is whether predicted energies recover the preferred transformation. Across 340 Li/Hong reaction groups per downstream seed, LARK ranked the true best site first in 95.98% of groups and second in every remaining group (Fig. 2e). Top-1 accuracy rose from 65.1% for energy gaps below 0.25 kcal mol^*−*1^ to 100% for gaps above 2 kcal mol^*−*1^. Thus, energy prediction translated into accurate site selection, with ranking errors concentrated among near-degenerate candidates.

We next tested whether reaction prediction remained useful with limited labelled data. With 10% of the original Li/Hong labels, LARK retained an *R*^2^ of 0.8231 (Supplementary Table S1). Learning curves for Zahrt and two Li/Hong settings showed improving prediction as labelled fractions increased (Fig. 2f). Between 5% and 70% labelled data, Zahrt *R*^2^ rose from 0.456 to 0.873; the expanded and original Li/Hong settings reached 0.981 and 0.977, respectively. These results establish transfer under limited supervision and show the additional value of task-specific labels.

Yield prediction extended the evaluation to an experimentally measured reaction outcome. On 576 held-out Suzuki high-throughput experimentation reactions, LARK achieved an *R*^2^ of 0.881 and an MAE of 6.24 percentage points, alongside yield-dependent residual bias (Fig. 2g). Across these benchmarks, the learned representations supported continuous outcomes and chemically meaningful choices between reactive sites, including settings with limited labelled data.

### Reaction representations support prediction across heterogeneous patent reactions

To extend the evaluation beyond individual chemical systems, we tested classification, condition inference and yield prediction on heterogeneous patent reaction records (Fig. 3 and Supplementary Table S1). On USPTO-Condition, exact condition-set accuracy increased from 30.15% at Top-1 to 52.75% at Top-15 across 68,073 Test reactions (Fig. 3a). At Top-15, LARK exceeded Reaction Graph, the strongest published comparator at the same candidate budget, by 0.95 percentage points[12–14]. Component-level decomposition exposed a remaining difficulty: solvent 2 and reagent 2 accuracies fell from 80.66% and 75.55% across all rows to 20.10% and 31.54% when recorded (Fig. 3b). Correctly predicting that no additional solvent or reagent was recorded boosted aggregate accuracy. Identifying these components when present remained substantially harder.

**Figure 3.**
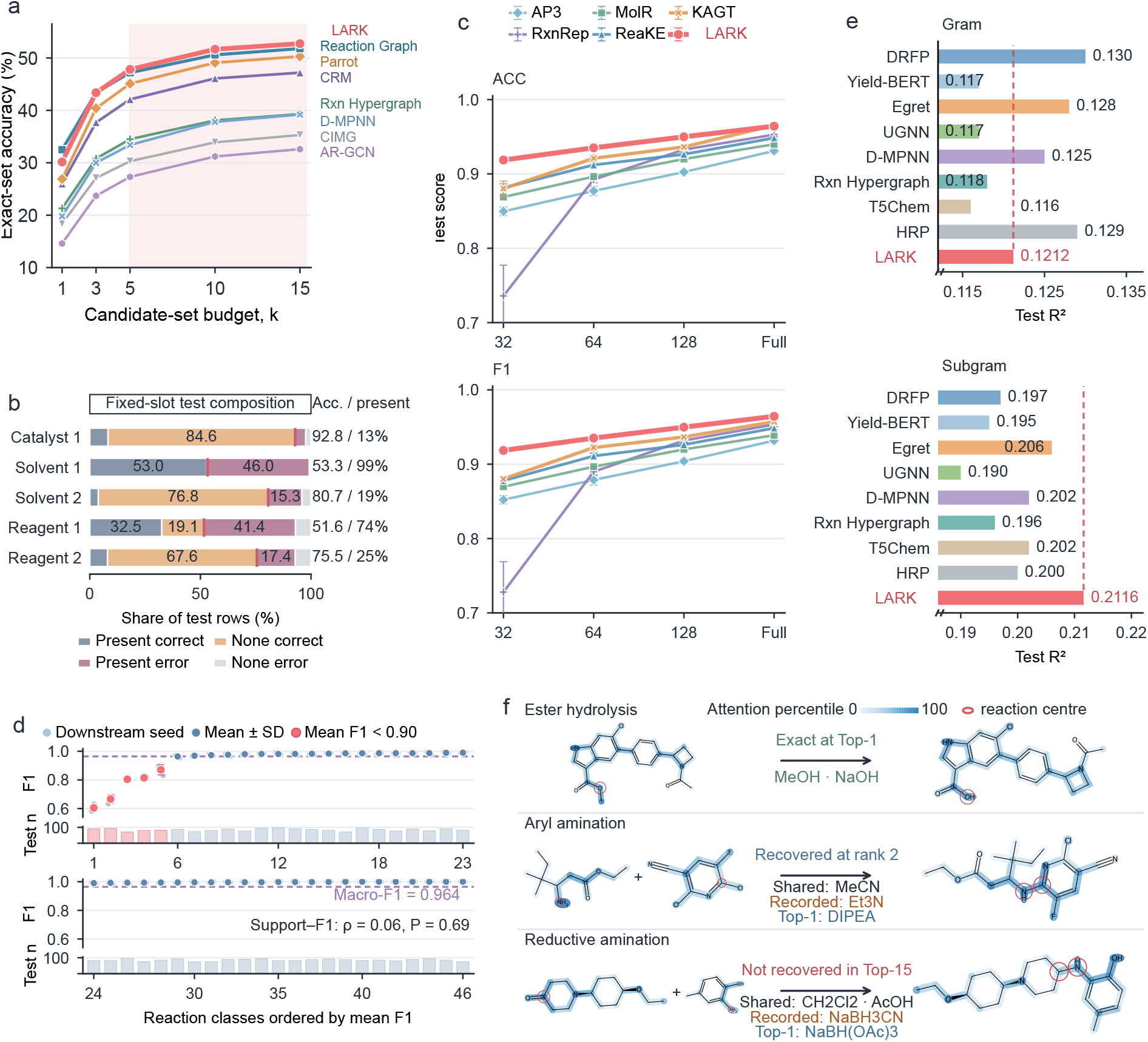
Patent-scale reaction classification, condition inference and yield prediction. **a**, Exact five-slot condition-set accuracy at increasing candidate budgets on 68,073 USPTO Test reactions. Curves are point estimates. **b**, Test-row composition for five condition slots. Each stacked row partitions present-correct, None-correct, present-error and None-error outcomes; the red marker gives all-row accuracy and the right label gives accuracy/present prevalence. **c**, Schneider reaction classification at 32, 64 and 128 labelled reactions per class and at full scale. LARK and published references report accuracy and macro-F1; points and error bars are mean sample *±* s.d. where repeats were available. **d**, Class-wise F1 over 46 classes. Pale points are three downstream seeds, dark points and whiskers are mean sample *±* s.d., the dashed line is overall macro-F1, red classes have mean F1 below 0.90, and the lower bars show fixed-Test support (*n* = 3,901). **e**, Gram and Subgram USPTO yield prediction. Values are Test *R*^2^ estimates and the dashed line marks LARK. **f**, Three fixed-slot outcomes: exact Top-1, recovery at rank 2, and absence from Top-15. Molecular colour encodes within-reaction encoder-attention percentile; red circles mark labelled reaction-centre atoms. Attention visualizes encoder-level localization.

Reaction classification tested whether the representations captured distinctions between transformation classes with sparse supervision. With 32 labelled examples per class, LARK exceeded the strongest displayed baselines by relative margins of 4.3% in accuracy and 4.4% in F1 (Fig. 3c)[4, 15]. At full scale, accuracy reached 0.9642 and F1 reached 0.9643; relative to KAGT, these were 0.1% lower and 0.8% higher, respectively. Performance extended across the class set, with 41 of 46 classes achieving mean F1 of at least 0.95 (Fig. 3d). Class Test support showed little association with class-wise F1 (Spearman *ρ* = 0.060, *P* = 0.691), indicating that class frequency alone did not account for the residual performance pattern.

Continuous yield prediction provided a more difficult transfer setting (Fig. 3e). LARK achieved an *R*^2^ of 0.121 on Gram reactions, 6.8% below DRFP, the strongest reported comparator. On Subgram reactions, its *R*^2^ of 0.212 was the highest reported value in this comparison, 2.7% above Egret[14, 16]. Both the low absolute scores and the reversed baseline ordering show that patent-yield prediction remains challenging and depends on the data regime.

Condition examples illustrated exact recovery, recovery at rank 2 and absence from Top-15, including ranking differences between chemically related reagents (Fig. 3f). Reaction-centre attention appeared in both successful and unsuccessful examples, showing that localized encoder signals can coexist with incorrect condition selection. Supplementary Fig. S1 further separates candidate coverage from ordering. Reaction pretraining thus supported classification, condition inference and yield regression within one framework, including accurate classification with sparse labels.

### Representations learned from reactions transfer to single-molecule properties

We next tested whether representations learned from reactions could also support property prediction for individual molecules. We compared LARK with 27 published methods across six MoleculeNet classification tasks (Supplementary Table S2). Among LARK and the 25 published methods reporting all six tasks, LARK achieved the highest mean AUROC of 0.779, compared with 0.755 for M2UMol (Fig. 4a)[3, 9]. Transfer also extended to continuous properties: LARK achieved the lowest reported RMSE among the methods compared for ESOL and Lipophilicity, although FreeSolv performance was weaker (Supplementary Table S3). The representations thus retained predictive information for both classification and regression from single-molecule inputs.

**Figure 4.**
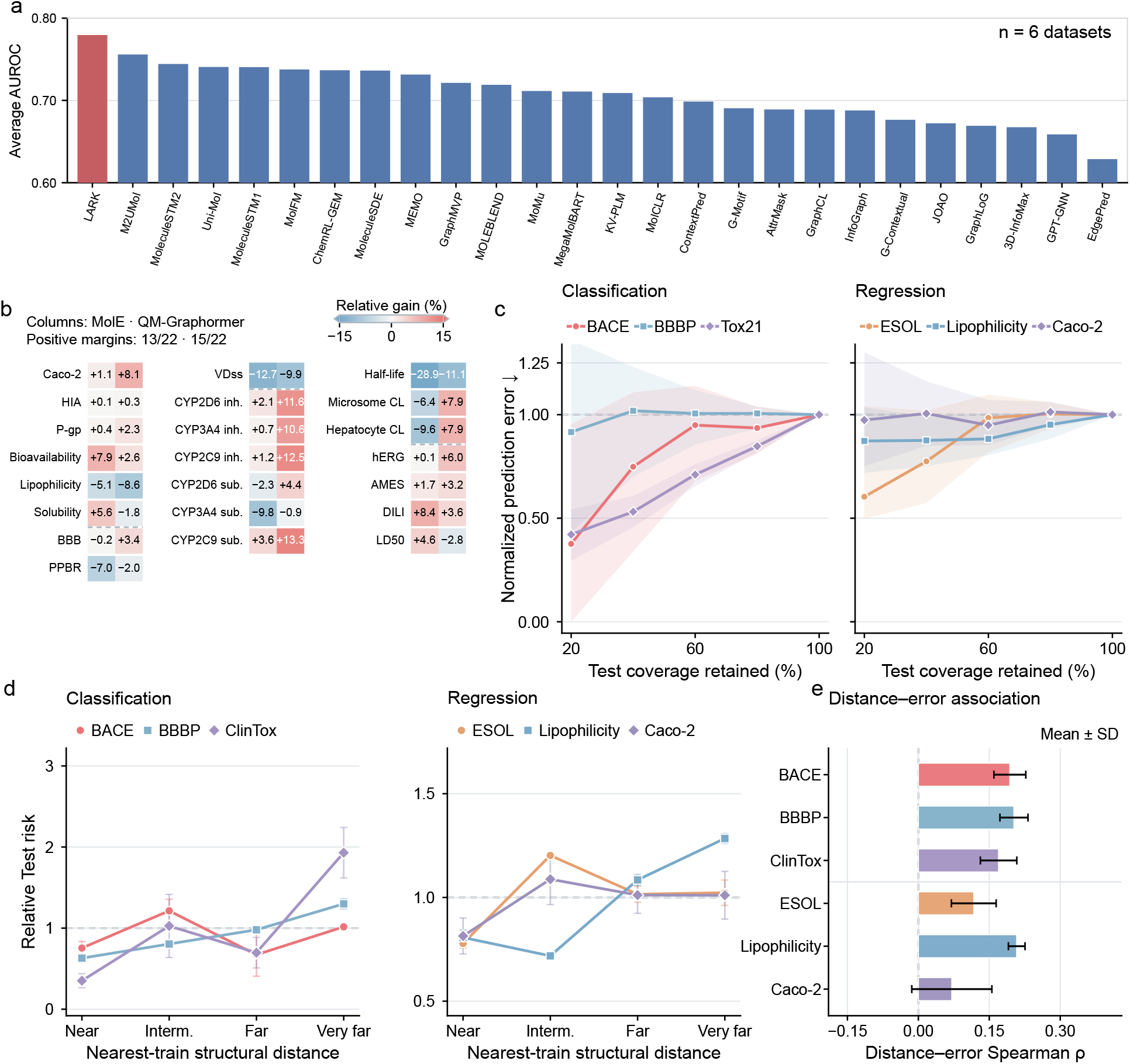
Molecular-property and ADMET transfer with reliability boundaries. **a**, Mean AUROC across BACE, BBBP, ClinTox, SIDER, Tox21 and ToxCast for LARK and the 25 published methods reporting all six tasks. Bars show the arithmetic mean of the six task AUROCs for each model. **b**, Direction-normalized relative mean difference between LARK and MolE (left cell) or QM-Graphormer (right cell) for 22 TDC ADMET tasks. Positive values favour LARK. Colour clipping applies at *±* 15%; printed values retain their original magnitude and each heterogeneous metric remains task specific. **c**, Normalized prediction error versus retained Test coverage for classification and regression tasks. Molecules are ordered by ensemble uncertainty; shading denotes 2,000-resample paired-molecule bootstrap 95% intervals. **d**, Relative Test risk across near, intermediate, far and very-far structural-distance strata. Distance is one minus maximum Morgan–Tanimoto similarity to downstream Train; cut-points were fixed from Validation. Points and error bars are mean *±* sample s.d. across downstream repeats. **e**, Mean *±* sample s.d. of the Spearman association between structural distance and per-molecule error (*n* = 3 downstream repeats, except Caco-2 *n* = 5).

We examined the breadth of this transfer across 22 TDC ADMET tasks. After accounting for metric direction, LARK outperformed MolE on 13 task means and QM-Graphormer on 15 (Fig. 4b and Supplementary Table S4)[2, 17]. Relative to MolE, Bioavailability AUROC was 7.9% higher, whereas half-life Spearman correlation was 28.9% lower. These results show that reaction pretraining yields molecular representations useful for predicting biological and pharmacokinetic endpoints, extending their utility beyond reaction outcomes.

We then asked whether reliability measures could identify where these predictions were most useful. Retaining the 20% least-uncertain Test molecules reduced normalized prediction error to 0.376 for BACE and 0.421 for Tox21 (Fig. 4c). The reduction persisted at broader coverage for Tox21 and was strongest in the most selective BACE subset. BBBP and the three regression tasks showed less consistent separation, making uncertainty informative for selective prediction in a subset of tasks.

Structural distance provided a second view of prediction reliability. In the very-far stratum, relative risk reached 1.30 for BBBP, 1.93 for ClinTox and 1.28 for Lipophilicity (Fig. 4d). Risk varied non-monotonically across distance strata for BACE, ESOL and Caco-2, with higher mean error in the intermediate stratum than in the neighbouring strata. These patterns indicate that structural similarity to the training set alone does not determine prediction difficulty: structurally closer molecules are not necessarily easier to predict than more distant ones. Continuous distance–error associations were positive across downstream repeats for BACE, BBBP, ClinTox and Lipophilicity, with mean Spearman correlations of 0.170–0.208 (Fig. 4e). Associations were weaker for ESOL and more variable for Caco-2. Uncertainty and structural distance therefore exposed different, task-specific patterns of error.

Single-molecule transfer establishes that the information learned through reaction pretraining remains useful beyond reaction prediction. The property and ADMET comparisons support this broader scope of chemical representation learning.

### Transformation supervision and reaction context improve transfer across reaction and molecular tasks

To identify which parts of the pretraining design supported molecular and reaction transfer, we compared eight variants differing in transformation supervision, directional views and reaction context (Fig. 5a). Scratch omitted pretraining. The single-view baseline A used masked-atom and fingerprint objectives; geometry yielded B, and adding ΔBE yielded C. D was the dual-view counterpart of B; adding ΔBE yielded E, consistency yielded F, and unmapped-component context completed LARK. Pretrained variants shared the corpus, architecture family, pretraining realization and training stage. A–C processed one directional view per reaction, whereas D–F and LARK processed both.

**Figure 5.**
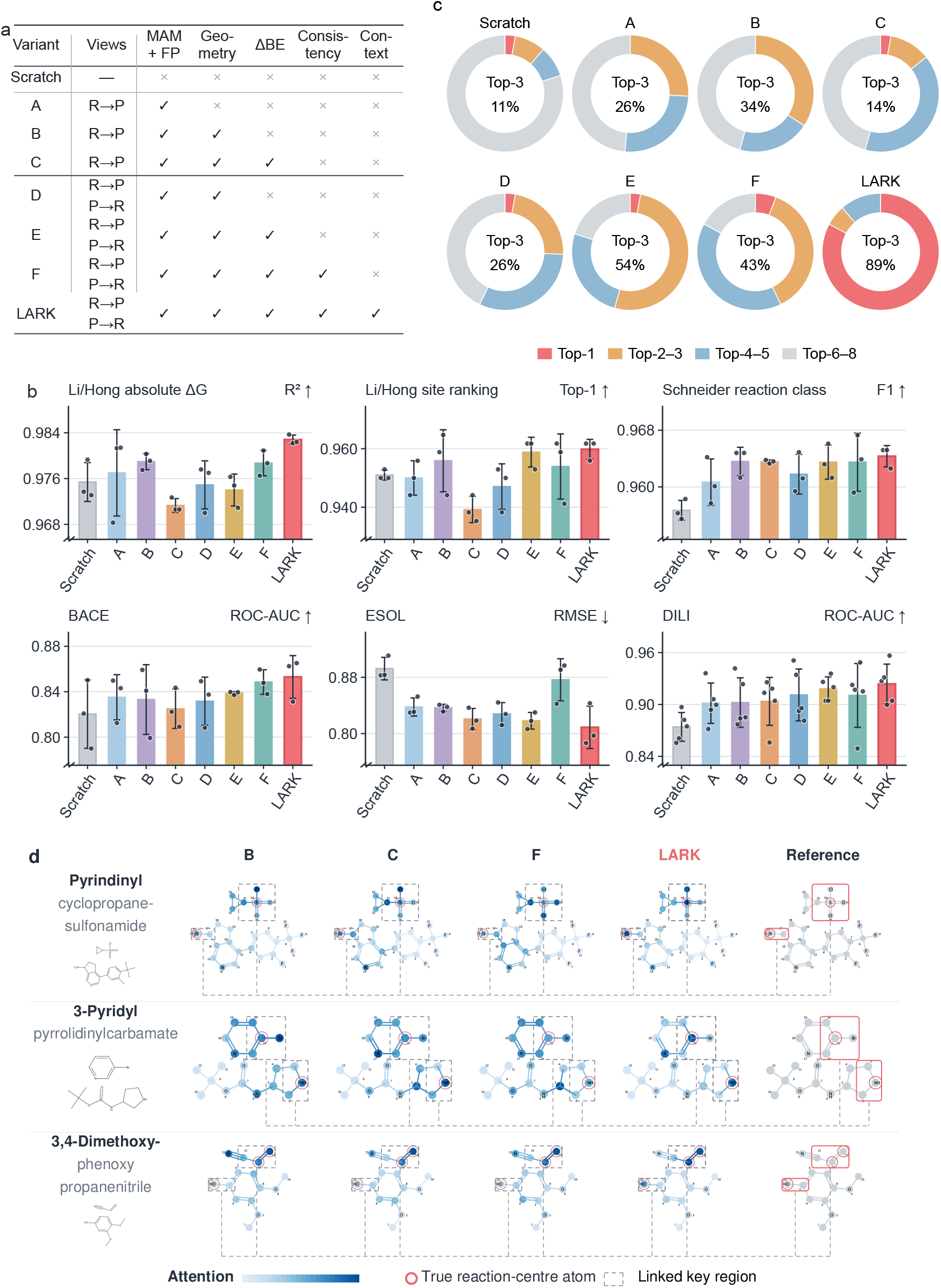
Matched ablations resolve task-dependent component contributions. **a**, Pre-specified Scratch and A–F controls alongside LARK. A–C use a single R *→* P view; D–F and LARK use dual R *→* P/P *→* R views. **b**, Six representative downstream tasks. Points are repeat values; bars and whiskers are mean *±* sample s.d. across three downstream repeats, except DILI (five frozen splits). Lower is better for ESOL RMSE; higher is better otherwise. All pretrained variants use one pretraining seed. **c**, Rank-tier composition across 35 tasks for which all eight models are available. Ranks are computed within task after respecting metric direction; the centre reports the fraction in the top three. **d**, Three reactions selected by fixed rules for complete LARK centre recovery, comparing B, C, F and LARK atom attention with the labelled reference. Attention is normalized within each model/reaction; dashed regions link corresponding substructures. Fig. 6a–c evaluates LARK localization and atom-state interventions across the larger analysis subset.

ΔBE supervision improved downstream performance across a range of reaction and molecular property tasks. Adding it to single-view geometry pretraining (B to C) improved 20 of 39 comparable task means, with reduced Li/Hong absolute-energy performance and nearly unchanged Schneider classification (Fig. 5b and Supplementary Table S5). In the dual-view configuration (D to E), ΔBE improved 26 of 39 task means, including seven of nine MoleculeNet tasks. With ΔBE supervision, adding the second view (C to E) improved 29 of the 35 completely matched tasks, including Li/Hong regression and ranking, BACE and ESOL. These contrasts support broader transfer gains from transformation supervision when learned through both directional views.

Consistency and reaction context further shaped transfer. Consistency (E to F) improved 23 of 39 comparable task means, including BACE and Li/Hong few-shot regression, but reduced ESOL performance. Adding unmapped context (F to LARK) improved 36 of 39 task means. This pattern supports the contribution of surrounding reaction components alongside local transformation targets.

The complete model ranked first on 29 of the 35 tasks with all eight variants available and within the top three on 31 (Fig. 5c). It also achieved the best mean on all six representative tasks in Fig. 5b. Relative to Scratch, BACE AUROC increased by 4.0%, ESOL RMSE decreased by 9.3% and DILI AUROC increased by 5.7%. The task-wide atlas and repeat variation are reported in Supplementary Fig. S2 and Supplementary Table S5.

Three reactions selected for complete LARK centre recovery illustrated how attention differed between configurations (Fig. 5d). In each, the top-*k* LARK atoms covered all *k* labelled centre atoms, whereas B, C and F placed at least one unchanged atom above a centre atom. Together with the downstream comparisons, these findings support combining transformation supervision with molecular structure, geometry and reaction context to improve transfer across reaction and molecular property tasks.

### Reaction-centre representations support bond–electron reconstruction

To extend the selected ablation examples to a larger reaction set, we tested whether LARK localized chemical changes and used highly attended atom states during reconstruction. The analysis used a fixed clean subset of 5,000 Test reactions evaluated in both directions (Fig. 6). We measured molecular-token-to-atom attention in the final molecular-encoder layer, averaged across heads. Attention was enriched at ΔBE-defined centre atoms above within-view prevalence, with mean average-precision enrichment of 0.335 for P → R and 0.453 for R → P (Fig. 6a). Predicted ΔBE magnitude localized these atoms more strongly, reaching 0.584 and 0.723, respectively. The rank-based combination did not improve on predicted changes alone. Thus, both internal attention and explicit reconstruction outputs carried spatial information about transformed atoms.

**Figure 6.**
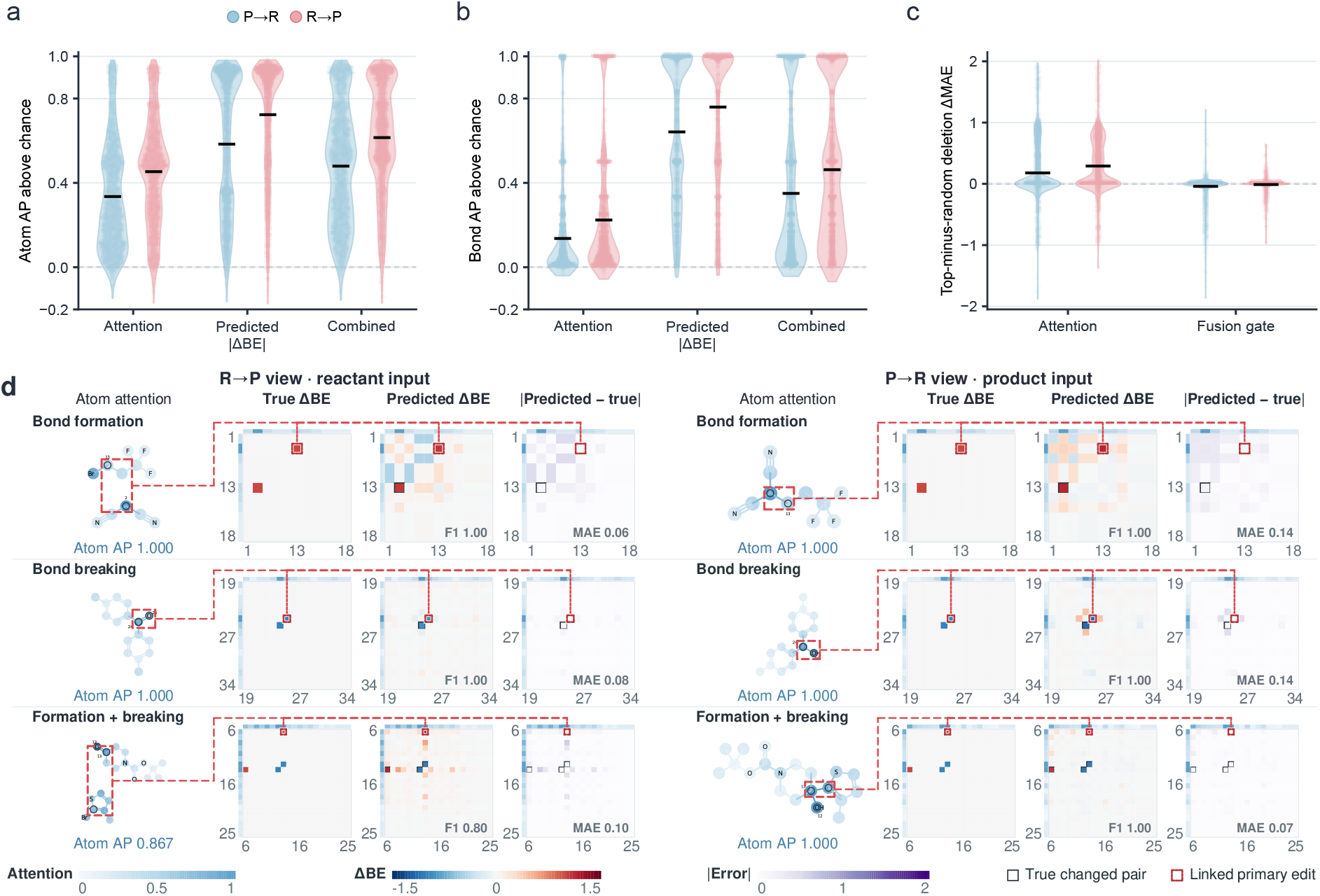
Reaction-centre information grounds directional representations. **a,b**, Per-reaction distributions of atom- and bond-level average-precision enrichment above the within-view reaction-centre prevalence for attention, predicted |ΔBE| and their combination. Pale points show all finite reaction values, violins show their distributions and black horizontal lines denote means. Panel a contains 4,368 finite reactions per direction and panel b contains 4,222. **c**, Per-reaction increase in changed-pair ΔBE MAE after suppressing the top-ranked atom representation relative to matched random atom-state suppression. Attention and fusion-gate rankings are shown for 4,222 finite reactions per direction; points, violins and black lines denote individual reactions, distributions and means, respectively. **d**, Three rule-selected successes spanning bond formation, bond breaking and mixed formation/breaking. For each direction, atom attention is linked to true, predicted and absolute-error ΔBE matrices; outlined cells are true changed pairs at |ΔBE| *≥* 0.5. Panels a–c provide the population distributions, and panel d illustrates them at molecular level.

Bond-level analysis showed the same ordering, with predicted |ΔBE| providing stronger localization than attention and both signals showing positive mean enrichment (Fig. 6b). Localization was stronger from reactant input at both atom and bond levels. Evaluation over the complete clean reaction collection further assessed bond formation, bond breaking and diagonal electron changes, with component-dependent reconstruction gains and cross-view agreement (Supplementary Fig. S3). These analyses link the learned signals to the locations and directional changes encoded by the reaction targets.

Matched interventions tested whether highly attended atom states contributed to reconstruction. Suppressing one highest-attention atom state per view increased MAE on true changed pairs by 0.178 in P→R and 0.290 in R→P beyond matched random suppression (Fig. 6c). Ranking atoms by fusion-gate magnitude did not produce the same additional disruption. Extended pathway interventions supported this distinction, while top-attended atoms varied across layers and attention heads (Supplementary Fig. S4). The localization and intervention results jointly indicate that highly attended atom states carry information used to reconstruct local bond–electron changes.

Three rule-selected successful reactions connected these population patterns to bond formation, bond cleavage and mixed edits (Fig. 6d). In both single-edit reactions, attention concentrated on the edited bond’s atoms, and predicted matrices recovered its location and sign with changed-pair F1 of 1.0 in both directions. In the mixed-edit case, P→R recovered both edits exactly; R→P recovered both but added one spurious changed pair, reducing F1 to 0.80. Intermediate and failure cases showed partial localization together with missed or additional matrix entries (Supplementary Fig. S5).

Together, spatial enrichment, matched interventions and signed edit recovery connect learned atom representations to the local bond–electron reorganization underlying chemical transformations. Alongside the transfer results, this evidence supports a framework that learns useful molecular and reaction representations while retaining an explicit connection to where chemical change occurs.

## Discussion

Chemical transformations provide supervision for learning how molecular structure relates to chemical behaviour. LARK brings this information into representation learning by combining observed reaction changes with molecular structure and geometry. The resulting representations support prediction at both molecular and reaction scales. This extends the scope of reaction pretraining: transformations can help organize representations of individual molecules as well as the reactions in which they participate.

Transfer to single-molecule properties is central to this interpretation. Learning within multicomponent reactions could favour information specific to reaction outcomes, yet the molecular encoder retains predictive value when reaction partners and context are absent from the downstream input. Together with transfer across reaction outcomes and condition inference, this finding supports chemical change as a useful complement to structural and geometric supervision. The framework thus connects two prediction settings through a shared pretraining design, with benefits extending beyond the reconstruction task used to learn the representations.

The reaction-centre analyses give this transfer framework a chemically interpretable basis. Attention and predicted changes localize to transformed atoms, and matched suppression shows that highly attended atom states carry information used in reconstruction. The intervention evidence strengthens the interpretation beyond attention localization alone[18]: learned atom states participate in recovering the local bond–electron reorganization that connects reactants and products. The ablations further support learning these changes through both directional views and incorporating surrounding reaction components. Together, these observations motivate a design in which local transformation supervision is embedded within molecular and reaction context, while structural and geometric objectives preserve information beyond the edited sites.

Transfer gains and prediction reliability vary across molecular targets and chemical distributions. Downstream component comparisons used one pretraining realization with different cumulative directional exposures; published benchmarks also differed in data partitions. Chemically, the supervision captures net bond and valence-electron changes between reactants and products, grounding the learned representations in the reorganization observed between reaction endpoints.

Future work can test this approach on unfamiliar reaction families and prospectively measured outcomes, and examine whether supervision from intermediates or reaction energetics improves transfer and chemical interpretation. The present results establish a practical route for integrating chemical change with molecular structure in pretraining, supporting chemical representations that remain useful across molecular properties and reaction behaviour.

## Methods

### Pretraining corpus, molecular records and data partitions

Pretraining reactions were prepared from the ORDerly release of the Open Reaction Database[19, 20]. Each record was decomposed into atom-mapped reactant components, atom-mapped product components and entirely unmapped components. The mapped components define the chemical transformation, whereas the unmapped molecules were retained as generic reaction context. Reaction sides were canonicalized after atom-map annotations had been removed, while stereochemistry and component multiplicity were preserved. Valid molecules were converted to attributed graphs and stored in indexed molecular records. An unmapped component that could not be parsed was omitted from the context set without discarding an otherwise valid mapped reaction.

A registry of downstream Validation and Test identities was frozen before the pretraining partitions were materialized. Each reaction in this registry was represented at three levels: the complete reaction, the complete product side and the largest-heavy-atom product core. Before pretraining, we excluded every reaction record whose complete-reaction, complete-product or largest-heavy-atom product-core identity matched an entry in the frozen downstream Validation or Test registry. The same identity definitions separated the clean internal Test partition from the candidate training pool. Additional input-side group filtering excluded candidates overlapping the internal Test partition. The final pretraining dataset comprised approximately 600,000 reactions across all three partitions: approximately 514,000 Train, 27,000 Validation and 60,000 clean Test reactions. The exposure audit screened approximately 555,000 candidate training records before the additional input-side filtering. The resulting partitions have zero overlap under these exact-reaction and mapped-core identities, while allowing chemically related molecules and scaffolds to occur in different partitions. The Validation partition was used for pretraining model selection, and the clean Test partition was reserved for the directional and reaction-centre analyses.

Downstream evaluation comprised 11 reaction and synthesis tasks, nine MoleculeNet tasks[9] and 22 ADMET tasks from Therapeutics Data Commons[10]. The split identity, target definition and primary metric were fixed separately for every task and then reused across all compared LARK variants. Missing labels in multi-task molecular datasets were masked during both optimization and metric calculation. Target transformations, class weights and any other label-dependent quantities were estimated from the downstream Train split only. The pretrained molecular encoder was fine-tuned on MoleculeNet and TDC tasks, using the fixed dataset partitions described in the Supplementary Methods.

### Molecular graphs and bond–electron change matrices

Each molecule was represented as a two-dimensional attributed graph with atoms as nodes, covalent bonds as edges and one learned molecular super-token. Atom attributes encoded element identity, formal charge, valence, hybridization, hydrogen count, chirality, aromaticity and ring membership, together with continuous mass, van der Waals radius and partial-charge features. Bond attributes described bond type, conjugation, ring membership, direction and stereochemistry. Discrete attributes were embedded and continuous values were expanded through learnable radial basis functions. The resulting channels were combined to form the initial atom states. Shortest-path distance and the bond features along each path supplied pair-specific attention biases. A molecular graph encoder jointly updated the atom states and the super-token, yielding an atom-level representation and one graph-level molecular representation from the same forward pass.

For an atom-mapped molecule, a bond–electron (BE) matrix was constructed on the coordinate system defined by its atom-map numbers. For atoms *i* and *j*, its entries were

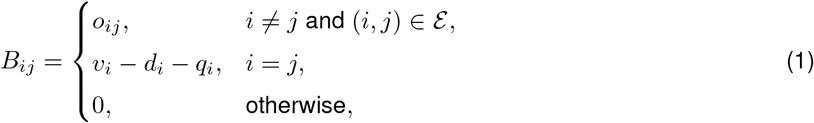

where ε is the set of covalent bonds, *o*_*ij*_ is the bond order, *v*_*i*_ is the outer-shell valence-electron count, *d*_*i*_ is the explicit valence and *q*_*i*_ is the formal charge. Bond orders were 1, 2 and 3 for single, double and triple bonds, respectively, and 1.5 for an aromatic bond. Component matrices on the same reaction side were padded to the common atom-map canvas and summed. This gave reactant- and product-side matrices, **B**_R_ and **B**_P_, with identical coordinates.

The canonical directional supervision target was

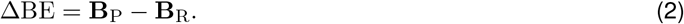

Thus, positive off-diagonal entries denote bond-order increases and negative entries denote bond-order decreases in the forward reaction direction; diagonal entries retain the associated atom-local electron redistribution. The matrix is symmetric, and a valid-canvas mask excludes padded coordinates. Its rows and columns index mapped-core atoms. Unmapped contextual molecules enter the reaction hierarchy as contextual inputs and modulate the encoded core representations.

### Dual-view reaction learning and hierarchical encoding

Each mapped reaction generated complementary reactant-to-product and product-to-reactant learning examples with shared parameters. In the reactant-to-product (R → P) view, the encoder received the mapped reactant core together with generic unmapped context and predicted +ΔBE. In the product-to-reactant (P → R) view, it received the mapped product core with the same context and predicted −ΔBE. The mapped reactant and product sides jointly defined these directional targets, while each encoder pass received the source-side mapped core for its direction. Together, the two directions associate each input core and its reaction context with a single canonical chemical change. Writing *G*_R_, *G*_P_ and *C* for the mapped reactant core, mapped product core and generic context, respectively, the two learning examples were

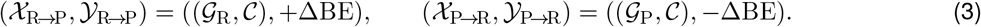

Encoding proceeded at three connected levels. First, the molecular graph encoder produced local atom states and one molecular token for every mapped or contextual component. Second, the role-aware reaction hypergraph Transformer (DirectedHyperGT) treated molecular components as nodes and each reaction as a hyperedge linking its components. Each directional input comprised the source-side mapped components and generic context, with the hyperedge represented by an additional reaction token. Direction entered through incidence-derived reactant, product and context role encodings. Within each input, full self-attention jointly updated the component and reaction tokens; only padding positions were masked.

Let *N* be the number of molecular components in a reaction, **h**_*m*_ the molecular token for component *m*, and *r*(*m*) ∈ {R, P, C} its role. The projected component state **u**_*m*_, normalized role encoding **p**_*m*_, component input 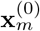, and reaction-token input 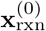 were

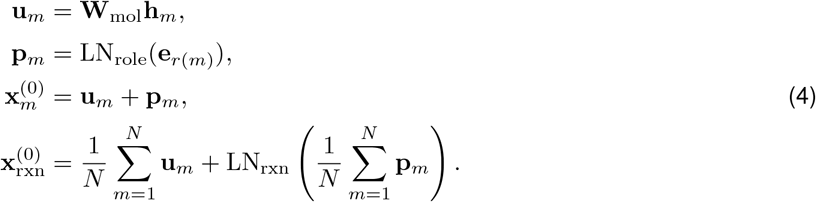

These states formed the sequence 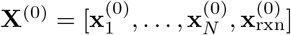

Each DirectedHyperGT layer used multi-head self-attention with padding masks, a sigmoid gate on the attended context, and residual, layer-normalized feed-forward updates. The full layer equations are provided in the Supplementary Methods. A final layer normalization returned contextualized component states and one reaction-level state.

Third, each contextualized molecular state was projected back to atom space and fused with its molecule-local atom states through a channel-wise gate,

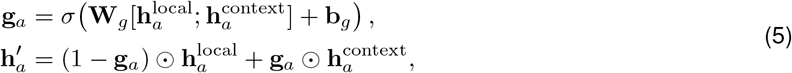

where brackets denote feature concatenation and *σ* is the sigmoid function. This feedback path allows reaction-level role and component information to reorganize atom-level representations before local pretraining targets are decoded. For the dual-view consistency objective, the contextualized mapped-component states were mean-pooled, with context components omitted from direct pooling. Context influenced the pooled core states through reaction-level attention; atom-state fusion supplied the local reconstruction heads.

The same hierarchy supplied representations for transfer. Single-molecule tasks used the molecular super-token. Reaction regression, ranking and classification tasks used the reaction-level representation or its task-specific role-aware extension. All pretraining heads were removed when the corresponding downstream head was attached.

### Multi-scale pretraining objectives

LARK used five multi-scale pretraining task types: (1) molecular fingerprint prediction, (2) masked atom modelling, (3) bond-angle prediction, (4) torsion-angle prediction and (5) directional ΔBE reconstruction. Molecular fingerprint prediction decoded a concatenation of Morgan, MACCS and RDKit fingerprints from the initial molecular token. Masked atom modelling reconstructed the identity of selected atoms from their surroundings, whereas the two local geometric tasks classified bond angles and torsion angles into discrete bins from the context-aware atom states. These five tasks provided supervision for mapped-core molecules at the molecule, atom, local-geometry and reaction-change levels. The masked atom modelling and fingerprint objectives were also applied to unmapped context molecules, providing direct structural supervision for their representations. A separate dual-view consistency term aligned the pooled mapped-core representations. The complete objective therefore comprised eight loss terms: five core-task losses, two context-task losses and one consistency loss.

Geometric labels were stored with the molecular records. The preprocessing routine used supplied coordinates when available; otherwise, for molecules with at most 400 atoms, it generated ten RDKit conformers, optimized them with MMFF and selected the lowest-energy conformer. Coordinate-generation failures and larger molecules used a two-dimensional fallback. Bond angles were assigned to 20 bins over [0, *π*], and torsions to 36 bins over [−*π, π*]; invalid geometric targets were masked. These labels supplied local geometric supervision.

For directional ΔBE reconstruction, the context-aware embeddings of mapped atoms were scattered onto the reaction canvas by atom-map identity. For every valid pair (*i, j*), a shared multilayer head operated on the two atom embeddings and returned a scalar change estimate. Averaging the two pair orders enforced a symmetric prediction,

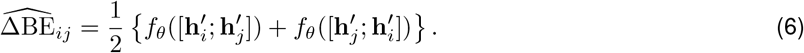

For reaction *r* and direction *v*, let *Y* ^(*r,v*)^ = *s*_*v*_ΔBE^(*r*)^, where *s*_R*→*P_ = +1 and 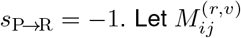 denote the valid-canvas mask and U = {(*i, j*) : *i* ≤ *j*} the upper triangle, including the diagonal. Writing B for the reaction views in a batch, reconstruction used the weighted mean over valid cells,

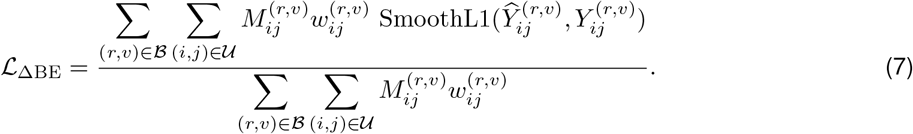

The cell weight was 10 when 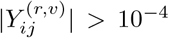 and 1 otherwise; SmoothL1 used transition parameter 1. This optimization threshold was distinct from the 0.5 threshold used to identify changed sites during evaluation. No target-derived BE pair feature was supplied to this head in the matched suite.

For dual-view variants, the mean-pooled mapped-core states from the two directional views were passed through a shared projector and normalized. For view *v* ∈ {R → P, P → R}, let M_*v*_ denote its mapped components and 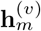 their contextualized molecular states. The projected view state and consistency loss were

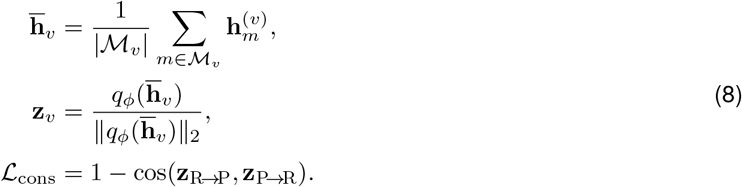

The core and context objectives were

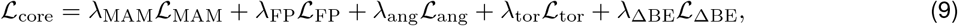

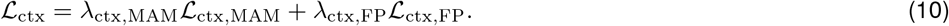

The complete pretraining objective was

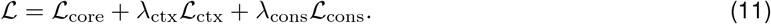

Enabled core-task coefficients were 1, as were the two context-task coefficients. LARK used *λ*_ctx_ = 1, and consistency-enabled variants used *λ*_cons_ = 0.1 with a 128-dimensional projector output. Consistency was averaged over paired reactions. In each ablation, loss terms associated with disabled components were removed from the training objective. During pretraining evaluation, molecular inputs were left unmasked, and directional ΔBE reconstruction was evaluated over all valid mapped-atom pairs.

### Matched ablations and model comparison

The pre-specified component suite comprised Scratch, variants A–F and LARK. Scratch initialized the downstream encoder randomly. A used one R → P view with masked-atom and fingerprint objectives. B added the bond-angle and torsion-angle objectives, and C added directional ΔBE reconstruction. D was the dual-view counterpart of B; E added directional ΔBE to D; F added mapped-core view consistency; and LARK added unmapped-component context. The planned contrasts evaluated geometry (A–B), ΔBE in a single-view model (B–C), dual views without ΔBE (B–D), dual views with ΔBE (C–E), ΔBE in the dual-view model (D–E), consistency (E–F) and generic context (F–LARK).

All pretrained variants used the same filtered corpus, architecture family, pretraining realization and training stage.

A–C processed one graph view per reaction, whereas D–F and LARK processed both directional views. Component effects were evaluated within each task because metric units and task demands differ across the downstream atlas.

The pretrained encoders were transferred to downstream tasks and fine-tuned with task-specific prediction heads. Task improvement counts compare mean metrics; repeats quantify downstream fitting or split variation from the shared pretraining realization. Repeat counts are reported with the corresponding results in the Supplementary Information.

### Downstream fine-tuning

For MoleculeNet and TDC tasks, the molecular encoder was fine-tuned with a task-specific prediction head. It produced one graph token per molecule, which was passed to a task-specific multilayer perceptron. Binary and multi-label tasks used masked, class-weighted binary cross-entropy; a label with a single class in an evaluation split was omitted from the macro average. Regression targets were standardized using Train-set statistics, optimized by mean-squared error in standardized units and transformed back to the original scale for evaluation. AUROC, average precision, MAE, RMSE and Spearman correlation were computed according to the task definition.

Li/Hong, Suzuki, Zahrt and USPTO yield regression attached a task head to the reaction representation. The head predicted a mean and log-variance and was optimized with a heteroscedastic Gaussian negative log-likelihood. Reported predictions used the mean. For Li/Hong site ranking, candidates belonging to the same reaction were kept in a fixed group and ordered by their predicted activation free energies; Top-1 accuracy indicates whether the experimentally preferred site received the lowest predicted energy. The few-shot series changed only the labelled fraction of the downstream Train set while retaining the Validation and Test identities of the corresponding few-shot series. Regression metrics were calculated in dataset-specific physical units before conversion to the displayed summary metric.

Schneider reaction classification used a multiclass head on the reaction representation and cross-entropy optimization. Test accuracy and macro-F1 were calculated over the fixed 46-class label space. Class-wise F1 was computed independently for each class before taking the unweighted macro average. Few-shot classification sampled the specified number of Train reactions per class and retained the full Validation and Test sets.

USPTO condition prediction represented catalyst 1, solvent 1, solvent 2, reagent 1 and reagent 2 as an ordered output sequence. Each slot was conditioned on the reaction representation and preceding slots, using recorded labels during teacher-forced training and predicted labels during inference. Missing components were represented by an explicit None class. Beam search produced complete candidate tuples, and exact-set accuracy measured whether the recorded tuple occurred among the Top-1, 3, 5, 10 or 15 candidates. Slot-level analyses separated all-row accuracy from accuracy among rows in which that component was recorded. Gram and Subgram yield tasks used the same reaction-transfer principle and reported Test *R*^2^.

### Selective prediction and structural-distance analysis

Reliability analyses used ensembles of the recorded downstream repeats for six representative molecular tasks. Classification predictions were averaged as probabilities, and predictive entropy of the mean probability defined molecular uncertainty. For regression, uncertainty was the sample standard deviation of the predictions across repeats. Molecules were ordered from lowest to highest uncertainty, the least-uncertain 20, 40, 60, 80 and 100% were retained, and prediction error was recomputed at each coverage. Classification error was measured by the Brier score, whereas regression retained the task’s MAE or RMSE definition. Each value was normalized by the corresponding error at 100% Test coverage.

Structural novelty was defined as one minus the maximum Tanimoto similarity between a Test molecule and any molecule in its downstream Train set, using radius-2 Morgan fingerprints. The 25th, 50th and 75th percentiles of Validation-to-Train distance fixed the near, intermediate, far and very-far strata before Test errors were inspected.

Within every task, stratum risk was divided by the full-Test risk. A complementary continuous analysis calculated Spearman’s correlation between nearest-Train distance and per-molecule prediction error for each downstream repeat.

### Reaction-centre attribution and deletion analysis

A true off-diagonal atom pair was labelled as changed when |ΔBE_*ij*_| ≥ 0.5. A changed atom either participated in such a pair or had an absolute diagonal change that met the same threshold. Positive and negative off-diagonal changes denoted bond formation or bond-order increase and bond breaking or bond-order decrease, respectively. Predictions from both inference directions were converted to the canonical product-minus-reactant orientation.

The population analyses used a fixed subset of 5,000 clean internal Test reactions evaluated in both directions. Atom attribution was the final molecular-encoder layer’s molecular-super-token-to-atom attention, averaged across attention heads. A candidate bond received the mean attention score of its two atoms. The atom-level ΔBE score was the largest absolute predicted pair change incident to that atom, including its diagonal entry.The bond-level score was 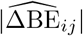. A combined score averaged the within-reaction percentile ranks of attention and predicted |ΔBE|. Average precision was calculated within each view and reported as enrichment above the corresponding changed-atom or changed-pair prevalence. Localization required both changed and unchanged candidates within a reaction view; undefined metrics were excluded from the corresponding summary. This yielded 4,368 finite atom-localization and 4,222 finite bond-localization values per direction.

Faithfulness was evaluated by matched representation deletion within each reaction. Under the same one-atom intervention budget, the highest-attention, a deterministically sampled random and the lowest-attention atom were selected. The selected atom’s final local state was set to zero after molecular encoding, while all other encoded states and the graph topology were retained. The altered states were then passed through the reaction hierarchy. The response was the increase in MAE over the true changed off-diagonal pairs, and top-minus-random effects were calculated within reaction before aggregation. The intervention summary contained 4,222 finite reactions per direction. Fusion-gate magnitude was the mean of the channel-wise atom-fusion gate and was evaluated using the same deletion procedure.

The molecular case studies were selected by fixed rules from successful bond-formation, bond-breaking and mixed-edit reactions. Atom-map identity linked each attribution map to the corresponding true, predicted and absolute-error ΔBE matrices. Separate ablation examples were selected where LARK recovered the complete labelled centre, and the same reaction identities were rendered for all partial variants.

### Statistical analysis

Selective-prediction intervals were estimated by paired molecule-level bootstrap resampling and reported as twosided 95% percentile intervals. The molecule or reaction was treated as the independent analysis unit. Multiple labels, atoms, bonds or beam candidates from the same unit were retained together during resampling or aggregation. The association between class support and class-wise F1 was evaluated using a two-sided Spearman rank-correlation test.

For task rankings, AUROC, average precision, accuracy, F1, *R*^2^ and Spearman correlation were ranked from high to low, whereas MAE and RMSE were ranked from low to high. Ties received average ranks. Relative differences were calculated as the metric difference divided by the absolute reference value and multiplied by For direction-normalized comparisons, the sign was reversed for lower-is-better metrics so that positive values favoured LARK. Absolute metric differences retain the metric’s units; accuracy differences expressed in percentage points are labelled explicitly.

## Data availability

Pretraining reactions were derived from the Open Reaction Database [https://github.com/open-reaction-database/ord-data] through the ORDerly release [https://figshare.com/articles/dataset/ORDerly_datasets/23502372]. The MoleculeNet datasets are publicly available through DeepChem [https://deepchem.readthedocs.io/en/latest/api_reference/moleculenet.html], and the ADMET datasets through the Therapeutics Data Commons ADMET Benchmark Group [https://tdcommons.ai/benchmark/admet_group/overview/].Public data for the reaction benchmarks are available from ChemSelML [https://github.com/Masker-Li/ChemSelML/tree/master/DataSet] for Li/Hong regioselectivity, the original study and its Supplementary Materials [https://doi.org/10.1126/science.aau5631] for Zahrt enantioselectivity, and Table 2 of the original Long and Ding study [https://doi.org/10.1002/1521-3773%2820010202%2940:3%3C544::AID-ANIE544%3E3.0.CO;2-8] for the Long/Ding enantiomeric-excess dataset. Public versions of the Suzuki– Miyaura and Buchwald–Hartwig yield datasets are available from rxn_yields [https://github.com/rxn4chemistry/rxn_yields/tree/master/data], and the Schneider reaction classification dataset from RXNRep [https://github.com/mjwen/rxnrep/tree/main/dataset/schneider]. USPTO-Condition is accessible on Hugging Face [https://huggingface.co/datasets/weidawang/USPTO_Condition], and the USPTO Gram and Subgram yield datasets are available through the rxn_yields data access page [https://github.com/rxn4chemistry/rxn_yields/blob/master/data/uspto/README.md].

## Code availability

All the codes are available at GitHub (https://github.com/KazeDog/LARK).

## Acknowledgements

The work was supported by the National Natural Science Foundation of China (No. 62322112) and the Science and Technology Development Fund of Macau (No. 0133/2024/RIB2).

## Notes

### Competing Interest Statement

The authors have declared no competing interest.

### Summary of Updates

Fixed some bugs and syntax issues.

## References

[1] Han Li, Ruotian Zhang, Yaosen Min, Dacheng Ma, Dan Zhao, and Jianyang Zeng. A knowledge-guided pre-training framework for improving molecular representation learning. Nature Communications, 14:7568, 2023.

[2] Oscar Méndez-Lucio, Christos A. Nicolaou, and Berton Earnshaw. MolE: a foundation model for molecular graphs using disentangled attention. Nature Communications, 15:9431, 2024.

[3] Zhankun Xiong, Ziyan Wang, Feng Huang, Minyao Qiu, Shuyan Fang, Liuqing Yang, Xionghui Zhou, Shichao Liu, Ping Zhang, and Wen Zhang. Multi-to-uni modal knowledge transfer pre-training for molecular representation learning. Nature Communications, 17:3797, 2026.

[4] Jiancong Xie, Yi Wang, Jiahua Rao, Shuangjia Zheng, and Yuedong Yang. Self-supervised contrastive molecular representation learning with a chemical synthesis knowledge graph. Journal of Chemical Information and Modeling, 64:1945–1954, 2024.

[5] Bo Qiang, Yiran Zhou, Yuheng Ding, Ningfeng Liu, Song Song, Liangren Zhang, Bo Huang, and Zhenming Liu. Bridging the gap between chemical reaction pretraining and conditional molecule generation with a unified model. Nature Machine Intelligence, 5:1476–1485, 2023.

[6] Jiacheng Xiong, Wei Zhang, Yinquan Wang, Jiatao Huang, Yuqi Shi, Mingyan Xu, Manjia Li, Zunyun Fu, Xiangtai Kong, Yitian Wang, Zhaoping Xiong, and Mingyue Zheng. Bridging chemistry and artificial intelligence by a reaction description language. Nature Machine Intelligence, 7:782–793, 2025.

[7] Li-Cheng Xu, Miao-Jiong Tang, Junyi An, Fenglei Cao, and Yuan Qi. A unified pre-trained deep learning framework for cross-task reaction performance prediction and synthesis planning. Nature Machine Intelligence, 7:1561–1571, 2025.

[8] Joonyoung F. Joung, Mun Hong Fong, Nicholas Casetti, Jordan P. Liles, Ne S. Dassanayake, and Connor W. Coley. Electron flow matching for generative reaction mechanism prediction. Nature, 645:115–123, 2025.

[9] Zhenqin Wu, Bharath Ramsundar, Evan N. Feinberg, Joseph Gomes, Caleb Geniesse, Aneesh S. Pappu, Karl Leswing, and Vijay Pande. MoleculeNet: a benchmark for molecular machine learning. Chemical Science, 9:513–530, 2018.

[10] Kexin Huang, Tianfan Fu, Wenhao Gao, Yue Zhao, Yusuf Roohani, Jure Leskovec, Connor W. Coley, Cao Xiao, Jimeng Sun, and Marinka Zitnik. Artificial intelligence foundation for therapeutic science. Nature Chemical Biology, 18:1033–1036, 2022.

[11] Xin Li, Shuo-Qing Zhang, Li-Cheng Xu, and Xin Hong. Predicting regioselectivity in radical C–H functionalization of heterocycles through machine learning. Angewandte Chemie International Edition, 59:13253–13259, 2020.

[12] Yu Zhang, Yang Han, Shuai Chen, Ruijie Yu, Xin Zhao, Xianbin Liu, Kaipeng Zeng, Mengdi Yu, Jidong Tian, Feng Zhu, Xiaokang Yang, Yaohui Jin, and Yanyan Xu. Large language models to accelerate organic chemistry synthesis. Nature Machine Intelligence, 7:1010–1022, 2025.

[13] Hanyu Gao, Thomas J. Struble, Connor W. Coley, Yuran Wang, William H. Green, and Klavs F. Jensen. Using machine learning to predict suitable conditions for organic reactions. ACS Central Science, 4:1465–1476, 2018.

[14] Yingzhao Jian, Yue Zhang, Ying Wei, Hehe Fan, and Yi Yang. Reaction graph: towards reaction-level modeling for chemical reactions with 3D structures. In Proceedings of the 42nd International Conference on Machine Learning, volume 267 of Proceedings of Machine Learning Research, pages 27426–27491, 2025.

[15] Jianbo Qiao, Kefei Li, Junru Jin, Ding Wang, Wenjia Gao, Shaoye Zhang, Shuan Liu, and Leyi Wei. Relation-aware pretraining and reaction center modeling for chemical reaction graph representation learning. Journal of Chemical Theory and Computation, 22(15):7774–7787, 2026.

[16] Daniel Probst, Philippe Schwaller, and Jean-Louis Reymond. Reaction classification and yield prediction using the differential reaction fingerprint DRFP. Digital Discovery, 1:91–97, 2022.

[17] Alessio Fallani, Ramil Nugmanov, Jose Arjona-Medina, Jörg Kurt Wegner, Alexandre Tkatchenko, and Kostiantyn Chernichenko. Pretraining graph transformers with atom-in-a-molecule quantum properties for improved ADMET modeling. Journal of Cheminformatics, 17:25, 2025.

[18] Sarthak Jain and Byron C. Wallace. Attention is not explanation. In Proceedings of the 2019 Conference of the North American Chapter of the Association for Computational Linguistics: Human Language Technologies, Volume 1 (Long and Short Papers), pages 3543–3556, 2019.

[19] Daniel S. Wigh, Joe Arrowsmith, Alexander Pomberger, Kobi C. Felton, and Alexei A. Lapkin. ORDerly: data sets and benchmarks for chemical reaction data. Journal of Chemical Information and Modeling, 64:3790–3798, 2024.

[20] Steven M. Kearnes, Michael R. Maser, Michael Wleklinski, et al. The open reaction database. Journal of the American Chemical Society, 143:18820–18826, 2021.

