## Supplementary information for "Reaction-aware hypergraph learning of chemical representations with LARK"

### A Experimental setup

#### A.1 Study design and evidence hierarchy

The evaluation was organized in three layers. First, LARK was assessed on reaction outcomes, reaction-site ranking, reaction classification, condition prediction and patent-scale yield prediction. Second, the same representation-learning framework was transferred to molecular-property and absorption, distribution, metabolism, excretion and toxicity (ADMET) tasks. Third, matched component ablations and reaction-centre analyses tested which parts of the pretraining design supported transfer and whether the learned signals were concentrated around the labelled chemical transformation.

The main text reports the representative comparisons needed for this evidence chain. The Supplementary Information records the datasets and split definitions, complete task matrix, secondary metrics, repeat-level results, candidate-set analyses, extended directional reconstruction and attention-intervention analyses, and representative intermediate and failure regimes. Supplementary Table S1 provides the published reaction- and synthesis-task comparisons underlying Figs. 2 and 3. Supplementary Tables S2 and S3 provide the task-resolved MoleculeNet classification and regression comparisons, and Supplementary Table S4 provides the absolute 22-task ADMET comparison underlying Fig. 4. Supplementary Table S5 provides the complete matched-ablation matrix as means and sample standard deviations across downstream repeats.

#### A.2 Pretraining corpus and molecular records

The matched ablation suite used the ORDerly downstream-holdout compact corpus. The final pretraining dataset comprised approximately 600,000 reactions, including approximately 514,000 Train, 27,000 Validation and 60,000 clean internal Test reactions. Before the compact corpus was materialized, a frozen registry of downstream Validation and Test identities was applied at the complete-reaction, complete-product and largest-heavy-atom product-core levels. Reaction sides were canonicalized after atom-map annotations had been removed, while stereochemistry and component multiplicity were retained. The same identity definitions separated the clean internal Test split from the candidate training pool.

Each record stored mapped reactant components, mapped product components and entirely unmapped components separately. Unmapped molecules supplied generic reaction context and were converted to the same attributed molecular graph representation as mapped molecules. Non-parsable context components were omitted while retaining an otherwise valid mapped reaction. Context components remained outside the atom-map canvas used to construct the directional $\Delta\mathrm{BE}$ target. Indexed compressed and uncompressed molecular stores contained the same graph tensors and therefore represented the same experimental condition.

The original ORDerly Train and Validation partitions were merged into a candidate pool of approximately 555,000 reactions. At training time, input-side group filtering removed approximately 14,000 candidates overlapping the clean internal Test partition, leaving approximately 541,000 reactions for training and validation. A deterministic subset of approximately 27,000 reactions was reserved for model selection, leaving approximately 514,000 reactions for parameter updates. The separate clean internal Test split of approximately 60,000 reactions was used only for locked directional and reaction-centre evaluation.

#### A.3 Downstream benchmark families

The task atlas comprised 11 reaction and synthesis tasks, nine MoleculeNet tasks and 22 Therapeutics Data Commons ADMET tasks. Reaction tasks covered activation-free-energy and selectivity regression, reaction-site ranking, yield prediction, few-shot transfer, reaction classification, reaction-condition prediction and Gram- and Subgram-scale patent yield prediction. The MoleculeNet panel comprised classification and regression benchmarks, whereas the ADMET panel covered absorption, distribution, metabolism, excretion and toxicity outcomes. Dataset names, the complete task list and evaluation metrics are reported in Supplementary Table S5.

Every model comparison within a task reused the same Train, Validation and Test identities. Missing labels in multi-task molecular datasets were masked during optimization and evaluation. Target transformations, class weights and other label-derived quantities were estimated from Train only. Few-shot experiments varied the labelled fraction or the number of labelled reactions per class while retaining the fixed Validation and Test sets.

#### A.4 Comparison and reporting

Published comparators were matched to the corresponding dataset and metric wherever the source study exposed a compatible protocol. Comparisons were made within tasks; metrics with different physical units or statistical meanings were not averaged into a common performance score. For the matched LARK suite, downstream heads, data partitions and training schedules were held fixed within each task. Seeded downstream tasks were generally repeated three times, while the TDC ADMET panel generally used five frozen split realizations. Means and sample standard deviations summarize those repeat units. Population-level reaction-centre analyses retain every finite reaction-level value and report arithmetic means; bootstrap intervals are reserved for the interval-based analyses specified below.

### B Supplementary Methods

#### B.1 Molecular graph and bond–electron representations

Each molecule was represented as an attributed graph with atoms as nodes, covalent bonds as edges and a learned molecular super-token. Atom attributes described element identity, formal charge, valence, hybridization, hydrogen count, chirality, aromaticity, ring membership and continuous physicochemical quantities. Bond attributes described bond type, conjugation, ring membership, direction and stereochemistry. Shortest-path distance and the bond features traversed by each path supplied pair-specific attention biases.

For an atom-mapped molecule, a bond–electron (BE) matrix was constructed on the coordinate system defined by atom-map numbers. Off-diagonal entries contained bond order and diagonal entries represented atom-local non-bonded valence-electron contributions. Component matrices on the same reaction side were padded to the common mapped-atom canvas and summed, giving reactant and product matrices, $\mathbf{B}_{\mathrm{R}}$ and $\mathbf{B}_{\mathrm{P}}$. The canonical directional target was

$$\Delta\mathrm{BE} = \mathbf{B}_{\mathrm{P}} - \mathbf{B}_{\mathrm{R}}. \tag{1}$$

Positive and negative off-diagonal values encode increases and decreases in bond order in the forward reaction direction; diagonal values retain atom-local electron redistribution. A valid-canvas mask excluded padded coordinates.

#### B.2 Dual-view reaction learning and context handling

Every mapped reaction could generate two complementary directional examples with shared model weights. In the reactant-to-product ($\mathrm{R} \to \mathrm{P}$) view, the encoder received the mapped reactant core and generic unmapped context and predicted $+\Delta\mathrm{BE}$. In the product-to-reactant ($\mathrm{P} \to \mathrm{R}$) view, the encoder received the mapped product core with the same context and predicted $-\Delta\mathrm{BE}$. The mapped reactant and product sides jointly defined these directional targets, while each encoder pass received the source-side mapped core for its direction.

Context molecules participated in hierarchical attention and gated atom–molecule fusion. The $\Delta\mathrm{BE}$ canvas indexed mapped-core atoms, and the dual-view consistency term was calculated from pooled mapped-core representations. Context components influenced these representations through the reaction hierarchy.

#### B.3 Hierarchical reaction encoder and pretraining objectives

The molecular graph encoder returned atom states and one molecular token for every mapped or contextual component. Molecular tokens were projected into reaction space, combined with learned reactant, product or context role embeddings and processed together with a reaction token by DirectedHyperGT. For a reaction containing $N$ molecular components, the reaction token was initialized from the mean projected component state. Its role encoding was obtained from the mean of the normalized component-role encodings.

The resulting sequence contained $N$ component tokens and one reaction token. Each DirectedHyperGT layer applied full multi-head self-attention, a token-wise sigmoid gate on the attended context, a residual layer-normalization update and a position-wise GELU feed-forward network. Padded component positions were masked during attention, and a final layer normalization returned the contextualized molecular states and the reaction-level state. For the layer equations below, $\mathbf{X}^{(l)}$ stacks component and reaction states, $j$ indexes heads and $l$ indexes layers.

For head $j$ in layer $l$, attention was computed as

$$\begin{aligned} &\mathbf{Q}_j^{(l)} = \mathbf{X}^{(l)}\mathbf{W}_{Q,j}^{(l)}, && \mathbf{K}_j^{(l)} = \mathbf{X}^{(l)}\mathbf{W}_{K,j}^{(l)}, && \mathbf{V}_j^{(l)} = \mathbf{X}^{(l)}\mathbf{W}_{V,j}^{(l)}, \\ &\mathbf{A}_j^{(l)} = \mathrm{softmax}\left(\frac{\mathbf{Q}_j^{(l)}\mathbf{K}_j^{(l)\top}}{\sqrt{d_j}} + \mathbf{M}_{\mathrm{pad}}\right), && \mathbf{C}^{(l)} = \mathrm{Concat}_j\left(\mathbf{A}_j^{(l)}\mathbf{V}_j^{(l)}\right). \end{aligned} \tag{2}$$

Here, $d_j$ is the head dimension and $\mathbf{M}_{\mathrm{pad}}$ masks padded component positions. A token-wise sigmoid gate modulated the attended context before the residual attention and feed-forward updates,

$$\begin{aligned} \mathbf{\Gamma}^{(l)} &= \sigma\left(\mathbf{X}^{(l)}\mathbf{W}_{\Gamma}^{(l)} + \mathbf{b}_{\Gamma}^{(l)}\right), \\ \widetilde{\mathbf{X}}^{(l)} &= \mathrm{LN}\Big[\mathbf{X}^{(l)} + \mathbf{W}_O^{(l)}\left(\mathbf{\Gamma}^{(l)} \odot \mathbf{C}^{(l)}\right)\Big], \\ \mathbf{X}^{(l+1)} &= \mathrm{LN}\Big[\widetilde{\mathbf{X}}^{(l)} + \mathbf{W}_2^{(l)}\,\mathrm{GELU}\left(\mathbf{W}_1^{(l)}\widetilde{\mathbf{X}}^{(l)} + \mathbf{b}_1^{(l)}\right) + \mathbf{b}_2^{(l)}\Big]. \end{aligned} \tag{3}$$

Dropout terms are omitted from the notation. A final layer normalization returned $N$ contextualized molecular states and one reaction-level state. The latter summarized the complete directional view.

The updated molecular state was projected back to atom space and combined with the molecule-local atom state through a learned channel-wise gate,

$$\mathbf{h}'_a = (1 - \mathbf{g}_a) \odot \mathbf{h}_a^{\mathrm{local}} + \mathbf{g}_a \odot \mathbf{h}_a^{\mathrm{context}}. \tag{4}$$

This hierarchy connects atom-, molecule- and reaction-level representations in a single forward pass.

LARK used five multi-scale pretraining task types: molecular fingerprint prediction, masked atom modelling, bond-angle prediction, torsion-angle prediction and directional local $\Delta\mathrm{BE}$ reconstruction. These five tasks supervised mapped-core molecules. Masked atom modelling and fingerprint prediction were also applied to unmapped context molecules, providing direct structural supervision for their representations. A separate dual-view consistency term aligned the pooled mapped-core representations. The complete objective therefore comprised eight loss terms: five core-task losses, two context-task losses and one consistency loss. The fingerprint target concatenated Morgan, MACCS and RDKit fingerprints. For $\Delta\mathrm{BE}$ reconstruction, context-aware mapped-atom states were scattered onto the atom-map canvas, and a shared pairwise decoder produced a symmetric matrix. Padded cells were masked and one triangular copy, including the diagonal, contributed to the reconstruction loss. For dual-view variants, projected mapped-core states were normalized and aligned by cosine distance. Condition vectors, target-derived BE pair features and the electron-conservation penalty were disabled in the matched suite.

#### B.4 Matched component ablations

The ablation matrix contained a Scratch reference trained on downstream tasks without pretraining, six partial variants and LARK. The component definitions were fixed before downstream comparison:

| **Model** | Views | Geometry | $\Delta\mathrm{BE}$ | Consistency | Context | Definition |
|---|---|---|---|---|---|---|
| **Scratch** | 0 | – | – | – | – | downstream encoder initialized without pretraining |
| **A** | 1 | – | – | – | – | $\mathrm{R}\to\mathrm{P}$ masked atom modelling and fingerprint prediction |
| **B** | 1 | ✓ | – | – | – | A plus bond-angle and torsion-angle objectives |
| **C** | 1 | ✓ | ✓ | – | – | B plus directional local $\Delta\mathrm{BE}$ supervision |
| **D** | 2 | ✓ | – | – | – | dual $\mathrm{R}\to\mathrm{P}$ and $\mathrm{P}\to\mathrm{R}$ counterpart of B |
| **E** | 2 | ✓ | ✓ | – | – | D plus directional local $\Delta\mathrm{BE}$ supervision |
| **F** | 2 | ✓ | ✓ | ✓ | – | E plus mapped-core view consistency |
| **LARK** | 2 | ✓ | ✓ | ✓ | ✓ | F plus generic unmapped-component context |

A, B and C used the $\mathrm{R}\to\mathrm{P}$ direction. D, E, F and LARK processed the dual views in the reporting order $\mathrm{R}\to\mathrm{P}$ followed by $\mathrm{P}\to\mathrm{R}$, with shared parameters across directions. The planned contrasts were A–B for geometry, B–C for single-view $\Delta\mathrm{BE}$, B–D for dual views without $\Delta\mathrm{BE}$, C–E for dual views with $\Delta\mathrm{BE}$, D–E for dual-view $\Delta\mathrm{BE}$, E–F for consistency and F–LARK for generic context. All variants used the same filtered corpus and pretraining stage. Complete-case ranks were calculated only for tasks containing Scratch, A–F and LARK.

For Supplementary Fig. S3, E(matched) used the same architecture and pretraining objectives as E, with half its cumulative directional graph-view exposure, matching that of C. The C-to-E(matched) comparison assessed dual-view training at equal cumulative view exposure; E(matched)-to-E assessed the effect of further training of the same configuration.

#### B.5 Downstream fine-tuning

##### B.5.1 Molecular-property and ADMET tasks

The transferred molecular encoder produced one graph token per molecule and a task-specific prediction head. Binary and multi-label tasks used masked, class-weighted binary cross-entropy. Let $y_{it} \in \{0,1\}$, $p_{it}$ and $m_{it} \in \{0,1\}$ denote the label, predicted probability and observed-label mask for molecule $i$ and task $t$, and let $\alpha_t$ be the positive-class weight estimated from the downstream Train split. The optimized loss was

$$\mathcal{L}_{\mathrm{BCE}} = -\frac{\sum_{i,t} m_{it}\left[\alpha_t y_{it}\log p_{it} + (1-y_{it})\log(1-p_{it})\right]}{\sum_{i,t} m_{it}\left[\alpha_t y_{it} + (1-y_{it})\right]}. \tag{5}$$

For regression task $t$, Train-set mean $\mu_t$ and standard deviation $s_t$ defined the standardized target $\widetilde{y}_{it} = (y_{it}-\mu_t)/s_t$. The prediction head was optimized by masked mean-squared error in standardized units,

$$\mathcal{L}_{\mathrm{MSE}} = \frac{\sum_{i,t} m_{it}(\widehat{\widetilde{y}}_{it} - \widetilde{y}_{it})^2}{\sum_{i,t} m_{it}}, \tag{6}$$

and predictions were transformed back to their original units for evaluation. The task-specific primary metrics were AUROC, average precision, MAE, RMSE or Spearman correlation. The encoder and prediction head were jointly fine-tuned unless the analysis was designated as a frozen-encoder probe.

##### B.5.2 Reaction regression, ranking and few-shot transfer

Li/Hong, Suzuki, Zahrt and USPTO yield regression used a heteroscedastic head on the reaction representation. The model returned a mean $\widehat{\mu}_i$ and log-variance $s_i = \log \widehat{\sigma}_i^2$ and minimized the Gaussian negative log-likelihood

$$\mathcal{L}_{\mathrm{NLL}} = \frac{1}{2N} \sum_{i=1}^{N} \left[\exp(-s_i)(y_i - \widehat{\mu}_i)^2 + s_i\right], \tag{7}$$

with the additive Gaussian constant omitted. Reported predictions used $\widehat{\mu}_i$. Li/Hong site-ranking candidates were grouped by reaction and ordered by their predicted activation free energies; Top-1 accuracy measured whether the preferred experimental site received the lowest predicted energy. Few-shot regression retained the Validation and Test identities within each few-shot series while varying the labelled Train fraction. These series used their own partitions and were not assumed to share Test identities with the full-data benchmark. Dataset-specific physical units were restored before the displayed metrics were calculated.

##### B.5.3 Reaction classification, condition prediction and patent yields

Schneider reaction classification used a multiclass head on the reaction representation and the standard cross-entropy loss

$$\mathcal{L}_{\mathrm{CE}} = -\frac{1}{N} \sum_{i=1}^{N} \log p_{i,y_i}, \tag{8}$$

where $p_{i,y_i}$ is the predicted probability assigned to the observed class. Test accuracy and F1 were calculated over the 46-class label space. Few-shot classification sampled the specified number of Train reactions per class while retaining the full fixed Validation and Test sets.

For USPTO condition prediction, catalyst, solvent and reagent components were represented as an ordered output sequence with an explicit `None` class for a missing component. Each slot was conditioned on the reaction representation and preceding slots, using recorded labels during teacher-forced training and predicted labels during inference. If $\mathcal{S}$ denotes the ordered condition slots and $\omega_s$ their specified loss weights, condition training minimized

$$\mathcal{L}_{\mathrm{cond}} = \frac{\sum_{s \in \mathcal{S}} \omega_s \mathcal{L}_{\mathrm{CE}}^{(s)}}{\sum_{s \in \mathcal{S}} \omega_s}, \tag{9}$$

where each slot loss was evaluated only on valid labels and the explicit `None` token was treated as an ordinary class. Beam search returned complete candidate tuples, and exact-set accuracy was evaluated at the specified Top-$k$ cutoffs. Gram and Subgram patent-yield tasks used the same reaction-transfer principle and reported Test $R^2$.

##### B.5.4 Standard evaluation metrics

AUROC was interpreted as the probability that a randomly selected positive example received a higher score than a randomly selected negative example, with half credit for a tie. The regression coefficient of determination and Spearman rank correlation were

$$\mathrm{AUROC} = \Pr(s^+ > s^-) + \tfrac{1}{2}\Pr(s^+ = s^-), \tag{10}$$

$$R^2 = 1 - \frac{\sum_i (y_i - \widehat{y}_i)^2}{\sum_i (y_i - \overline{y})^2}, \tag{11}$$

$$\rho_{\mathrm{S}} = \mathrm{corr}(\mathrm{rank}(y), \mathrm{rank}(\widehat{y})). \tag{12}$$

Accuracy, F1, MAE and RMSE followed their standard definitions. For multi-label classification, metrics were calculated independently for each evaluable label and then macro-averaged; labels containing only one class in an evaluation split were omitted.

#### B.6 Directional $\Delta\mathrm{BE}$ and reaction-centre evaluation

Predictions from both directions were converted to the canonical product-minus-reactant orientation before reporting. A true off-diagonal pair was labelled as changed when its absolute $\Delta\mathrm{BE}$ reached the fixed change threshold. A changed atom either participated in such a pair or had an absolute diagonal change that reached the same threshold. Formation, breaking and diagonal-electron changes were retained as separate strata.

The locked directional evaluation used the complete clean internal Test split of approximately 60 thousand reactions, with the two directions matched by reaction identity. Fixed-threshold precision, recall and F1, Top-1/3/5 reaction-centre localization, electron-conservation error and edit-family probes were calculated on the audited common target universe. Frozen representation probes used fixed Train, Validation and Test reaction samples, with probe selection performed on Validation.

#### B.7 Atom attention, enrichment and matched deletion

Atom attribution was defined as molecular-super-token-to-atom attention from the final molecular-encoder layer, averaged across attention heads. Changed-atom enrichment was quantified by AUROC and average precision. Fusion-gate magnitude was the channel mean of each atom's fusion gate. Reaction-level molecular attention was analysed as a distinct internal signal.

For localization, the attention score of a candidate atom was its head-averaged attribution, and the attention score of a candidate bond was the arithmetic mean of its two endpoint-atom scores. The atom-level $\Delta\mathrm{BE}$ score was the largest absolute predicted change involving that atom, including its diagonal entry; the bond-level score was the absolute predicted change for that atom pair. The combined score was the arithmetic mean of the within-reaction percentile ranks of attention and $\Delta\mathrm{BE}$. Let $y_{(k)}$ denote the binary changed-site label at rank $k$, after sorting $n$ candidates by decreasing score, and let $P = \sum_{k=1}^{n} y_{(k)}$. Average precision was

$$\mathrm{AP} = \frac{1}{P}\sum_{k=1}^{n}\left(\frac{1}{k}\sum_{\ell=1}^{k} y_{(\ell)}\right) y_{(k)}. \tag{13}$$

Using the within-view positive prevalence $\pi = P/n$ as the reference, AP enrichment was defined as $\mathrm{AP}-\pi$; AUROC enrichment was $\mathrm{AUROC}-0.5$. Metrics were calculated within each reaction-view candidate set and then summarized across reactions. Localization summaries required both changed and unchanged candidates; candidate sets lacking either label were excluded. This yielded 4,368 finite atom-localization and 4,222 finite bond-localization values per direction in Fig. 6. The matched changed-pair deletion analysis retained 4,222 finite reactions per direction.

Faithfulness used matched within-reaction interventions. One top-attended, one deterministically selected random and one low-attention atom were chosen under the same one-atom intervention budget. The selected atom's final local representation was set to zero while the molecular super-token, graph topology and all other states were retained. The primary response was the change in reconstruction error on true changed atom pairs. Complete gate-off and context-off interventions separately removed the atom–molecule fusion contribution and the unmapped-context pathway.

The population analysis used a fixed subset of 5,000 clean Test reactions, yielding 10,000 directional views. Layer, head, direction and token-order comparisons measured the stability of the attribution pattern. Case visualizations linked atoms to the true, predicted and absolute-error $\Delta\mathrm{BE}$ matrices by atom-map identity.

Layer agreement measured whether the first and last molecular-encoder layers selected the same top-attended atom after averaging over heads. Head consensus measured the fraction of final-layer heads selecting the same top-attended atom as the head-averaged attention.

#### B.8 Selective prediction and structural distance

Selective prediction ranked Test molecules by uncertainty estimated from an ensemble of downstream repeats. Classification uncertainty used predictive entropy, whereas regression uncertainty used dispersion across predictions. Error was recomputed at increasing retained coverage and normalized by the full-Test error.

For $K$ downstream repeats, the ensemble probability or regression prediction was $\bar{p}_{it} = K^{-1}\sum_{k=1}^{K} p_{it}^{(k)}$ or $\bar{y}_i = K^{-1}\sum_{k=1}^{K} \hat{y}_i^{(k)}$, respectively. For a classification molecule with $T_i$ evaluable labels, uncertainty was mean binary predictive entropy,

$$u_i^{\mathrm{cls}} = -\frac{1}{T_i}\sum_{t=1}^{T_i}\left[\bar{p}_{it}\log\bar{p}_{it} + (1-\bar{p}_{it})\log(1-\bar{p}_{it})\right], \tag{14}$$

whereas regression uncertainty was the sample standard deviation across repeat predictions,

$$u_i^{\mathrm{reg}} = \sqrt{\frac{1}{K-1}\sum_{k=1}^{K}\left(\hat{y}_i^{(k)} - \bar{y}_i\right)^2}. \tag{15}$$

At coverage $c \in \{0.2, 0.4, 0.6, 0.8, 1.0\}$, the subset $\mathcal{S}_c$ contained the least-uncertain $\lceil cN \rceil$ Test molecules. The displayed normalized prediction error was

$$\mathrm{NPE}(c) = \frac{\mathcal{R}(\mathcal{S}_c)}{\mathcal{R}(\mathcal{S}_1)}, \tag{16}$$

where $\mathcal{R}$ was the Brier error for classification and the task-specific MAE or RMSE for regression.

Structural novelty was one minus the maximum radius-2 Morgan-fingerprint Tanimoto similarity between a Test molecule and the downstream Train set. Specifically, for radius-2 Morgan fingerprint $F_2$,

$$d(x,\mathcal{T}) = 1 - \max_{z\in\mathcal{T}} \mathrm{Tanimoto}\left(F_2(x), F_2(z)\right). \tag{17}$$

Validation-to-Train distance quartiles fixed the near, intermediate, far and very-far strata before Test analysis. Within a stratum $g$, normalized prediction error was $\mathcal{R}(g)/\mathcal{R}(\mathrm{Test})$, using the same task-specific risk as in the coverage analysis.

#### B.9 Candidate-set and multi-answer analysis

Suzuki and Buchwald high-throughput experimentation (HTE) condition recommendation was evaluated inside experimentally measured candidate pools. Under the primary validity rule, a candidate was valid when its yield was within five percentage points of the best measured candidate in the same reaction group. Within-ten-point and within-group top-decile definitions supplied sensitivity analyses. Repeated split/model observations were first averaged within each of the 15 unique reaction groups and then resampled by reaction group.

The residual-corrected comparator added a component-based correction to the fixed LARK yield prediction. A factorization machine learned linear and pairwise interaction terms from categorical reactant, product and role-labelled condition features, including component-set identities. Its feature vocabulary and residual targets, measured yield minus LARK prediction, were constructed from Train data only. The correction minimized squared error on standardized training residuals; factor rank, regularization and checkpoint selection used Validation $R^2$. After reversing the residual standardization, the correction was added to the LARK prediction and clipped to the 0–100% yield range. Test yields were used to evaluate predictions and define valid candidates, but not to fit or select the correction. This post-training comparator was evaluated only in the supplementary HTE candidate-ranking analysis.

Forward prediction and retrosynthesis contained one recorded reference per reaction. Their candidate archives were decomposed into recorded-reference Hit@1, ranking miss and candidate-generation miss. Forward analysis used Beam-20 candidates and retrosynthesis used $K = 200$ candidates. The HTE set-valued estimand and the USPTO recorded-reference estimand were reported separately.

##### B.10 Statistical analysis

Downstream repeat summaries used the arithmetic mean and sample standard deviation,

$$s = \sqrt{\frac{1}{n-1}\sum_{i=1}^{n}(x_i - \bar{x})^2}, \tag{18}$$

where the independent repeat was a downstream training seed or frozen split identity. Reaction-centre enrichment and matched deletion were summarized from all finite per-reaction observations, with directional views linked by reaction identity in the source data. HTE and candidate-decomposition intervals used two-sided percentile bootstrap confidence intervals. The resampling unit was a molecule, reaction or unique reaction group according to the interval-based analysis, with labels and candidate rows belonging to the same unit kept together. Task ranks were computed after applying the higher- or lower-is-better direction of each metric. Selective-prediction and structural-distance intervals used molecule-level resampling and recomputed the retained subset or distance-stratum risk inside each bootstrap sample. HTE intervals resampled unique reaction groups, keeping all candidates and validity labels from a group together.

#### C Supplementary Results and Discussion

##### C.1 Candidate-set analysis separates coverage from ranking

To distinguish errors in candidate coverage from errors in ordering, we examined whether an acceptable answer was available in each candidate list and whether it was ranked first. The HTE analysis allowed multiple conditions to be valid according to measured yields; the forward and retrosynthesis analyses evaluated recovery of the single recorded answer.

Under the primary validity rule, uncorrected LARK achieved any-valid Hit@1/5 of 80.00/95.56% for Suzuki and 70.00/96.67% for Buchwald. Residual correction yielded 82.22/95.56% and 83.33/98.33%, respectively (Supplementary Fig. S1a,b). The sensitivity definitions retained the same qualitative separation between Top-1 selection and broader candidate coverage (Supplementary Fig. S1d). Thus, a valid condition was usually present within the short candidate list even when the first-ranked choice was incorrect, identifying candidate ordering as the main opportunity for improving HTE condition selection.

For Forward Beam-20, recorded-product coverage and Top-1 recovery reached 96.44% and 86.33%, partitioning the remaining reactions into 10.11% ranking misses and 3.56% generation misses. Retrosynthesis $K = 200$ covered the recorded precursor set for 74.78% of reactions; listwise reranking increased Top-1 recovery from 10.86% to 21.65% and left 53.13% ranking misses and 25.22% generation misses (Supplementary Fig. S1c). Ranking misses exceeded generation misses in both archives and accounted for most retrosynthesis errors. These decompositions identify existing candidate coverage as an opportunity for better selection, while the remaining generation misses require improvements to the candidate set itself.

##### C.2 Task-wide ablations reveal task-dependent component contributions

The task-wide atlas tests whether the representative gains in Fig. 5 extend across reaction, MoleculeNet and ADMET tasks (Supplementary Fig. S2). It contains 42 tasks, of which 35 have results for all eight variants. LARK ranked first on 29 of these complete tasks, although several tasks favoured a partial variant. Pairwise contrasts use all tasks available for the two variants being compared and can therefore include more tasks than the complete-case ranking.

Transformation supervision produced broader gains in the dual-view configuration than in the single-view configuration. Adding $\Delta\mathrm{BE}$ to dual-view pretraining (D to E) improved six of eight reaction tasks, seven of nine MoleculeNet tasks and 13 of 22 ADMET tasks. Adding the second view with $\Delta\mathrm{BE}$ supervision (C to E) improved 30 of 39 available task means and reduced nine; 29 of those improvements belonged to the 35-task complete subset used in Fig. 5c. Consistency (E to F) improved five of eight reaction tasks, four of nine MoleculeNet tasks and 14 of 22 ADMET tasks. Its effects included improved BACE performance but reduced ESOL performance, illustrating differences between endpoints within the same task family.

Unmapped reaction context produced the most widespread improvement in these pairwise contrasts, improving 36 of 39 task means when added to F. The atlas therefore supports combining transformation supervision and reaction context for broad transfer, while showing where partial configurations remain competitive. Counts refer to differences in task means, not statistical significance; Supplementary Table S5 reports the absolute means and sample standard deviations underlying the normalized atlas.

#### C.3 Directional reconstruction and atom-state interventions support the functional role of reaction-centre representations

To complement the population-level localization and intervention analyses in Fig. 6, we evaluated directional reconstruction across the complete clean internal Test set and separated bond formation, bond breaking and diagonal electron changes (Supplementary Fig. S3). Dual-view training, additional optimization, consistency and context affected these components differently. LARK showed the highest cross-view full-matrix cosine similarity and signed agreement, whereas changed-pair overlap remained comparable across the dual-view variants. Reconstruction was easier from reactant input than from product input. These results extend the assessment to the full clean reaction collection and show how reconstruction depends on both the type of chemical change and the available input direction.

Pathway and atom-state interventions tested their contributions to reconstruction (Supplementary Fig. S4). Attention was enriched at labelled changed atoms, and suppressing highly attended atom states disrupted changed-pair reconstruction more strongly than suppressing random or low-attention states. Removing the fusion or context pathway also increased reconstruction error, supporting their contribution to the prediction. However, fusion-gate magnitude did not provide the same centre-localization or atom-importance ranking signal. The interventions thus support a functional contribution from reaction-centre-focused atom states and from the pathways that integrate molecular and reaction context. Top-attended atoms varied across layers and attention heads (Supplementary Fig. S4f), distinguishing the enrichment of final-layer, head-averaged attention from agreement among individual internal components.

Intermediate and failure examples illustrated where localization and edit recovery became incomplete (Supplementary Fig. S5). Intermediate examples retained partial centre attention or edit recovery, whereas failure examples combined diffuse attention with missed or spurious matrix entries. In these selected cases, considered alongside the successes in Fig. 6d, clearer centre localization coincided with more complete directional matrix recovery.

### D Supplementary Figures

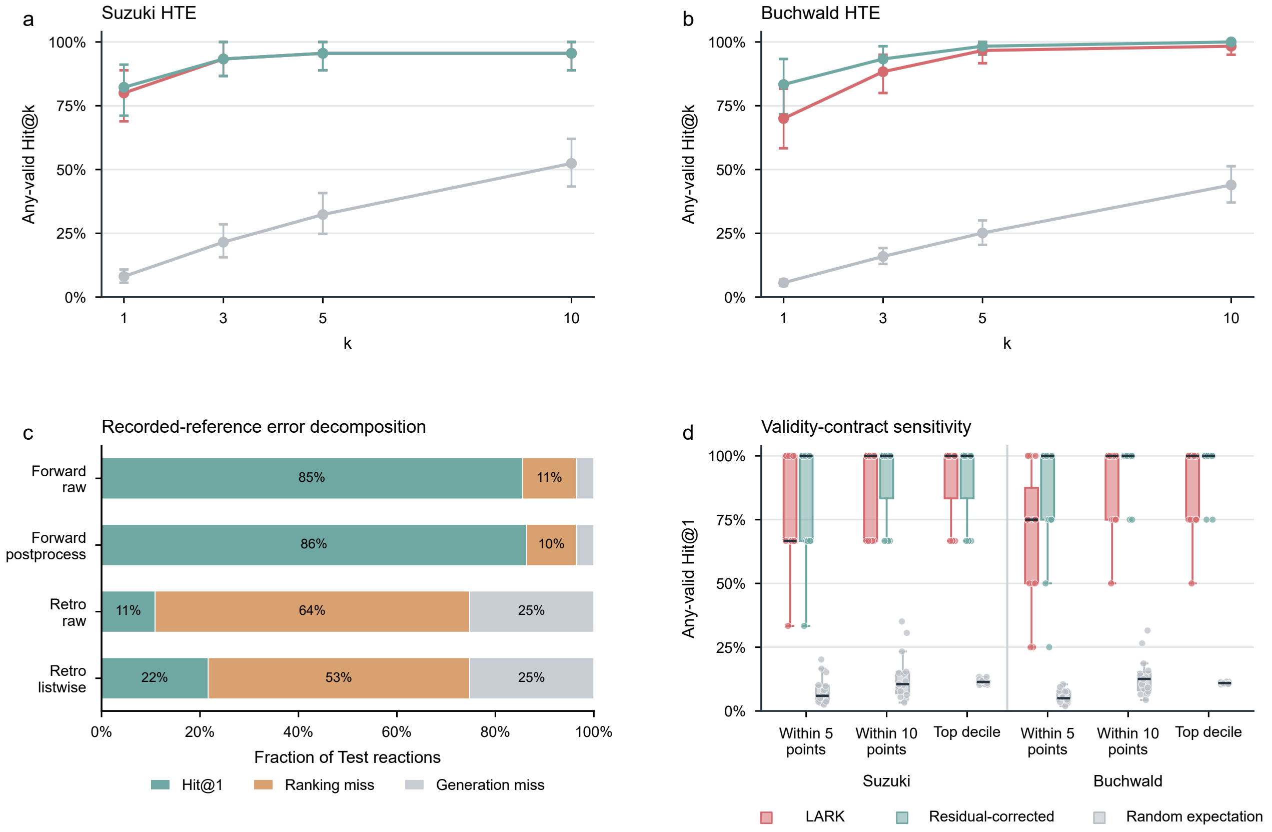

**Figure S1. Condition ranking and candidate coverage. a,b**, Any-valid Hit@$k$ for Suzuki and Buchwald HTE candidate pools under the primary validity rule (measured yield within five percentage points of the best candidate in the same reaction group). Points are macro means over 15 unique reaction groups; repeated split/model observations were averaged within group. Error bars are 2,000-resample reaction-group bootstrap 95% confidence intervals. **c**, Decomposition of Test reactions into recorded-reference Hit@1, ranking miss and candidate-generation miss for Forward Beam-20 and retrosynthesis $K = 200$. The three stacked fractions sum to one within each system. **d**, Any-valid Hit@1 under the primary within-five-point rule and the within-ten-point and top-decile sensitivity definitions. Each box summarizes 15 reaction groups after repeated split/model observations were averaged within group. Centre lines denote medians, boxes the interquartile range and whiskers extend to the most extreme value within 1.5 times the interquartile range; points show all individual reaction groups. In **a,b,d**, LARK denotes uncorrected yield predictions; Residual-corrected adds a component-based factorization-machine correction fitted on Train and selected on Validation, as described in Supplementary Methods B.9.

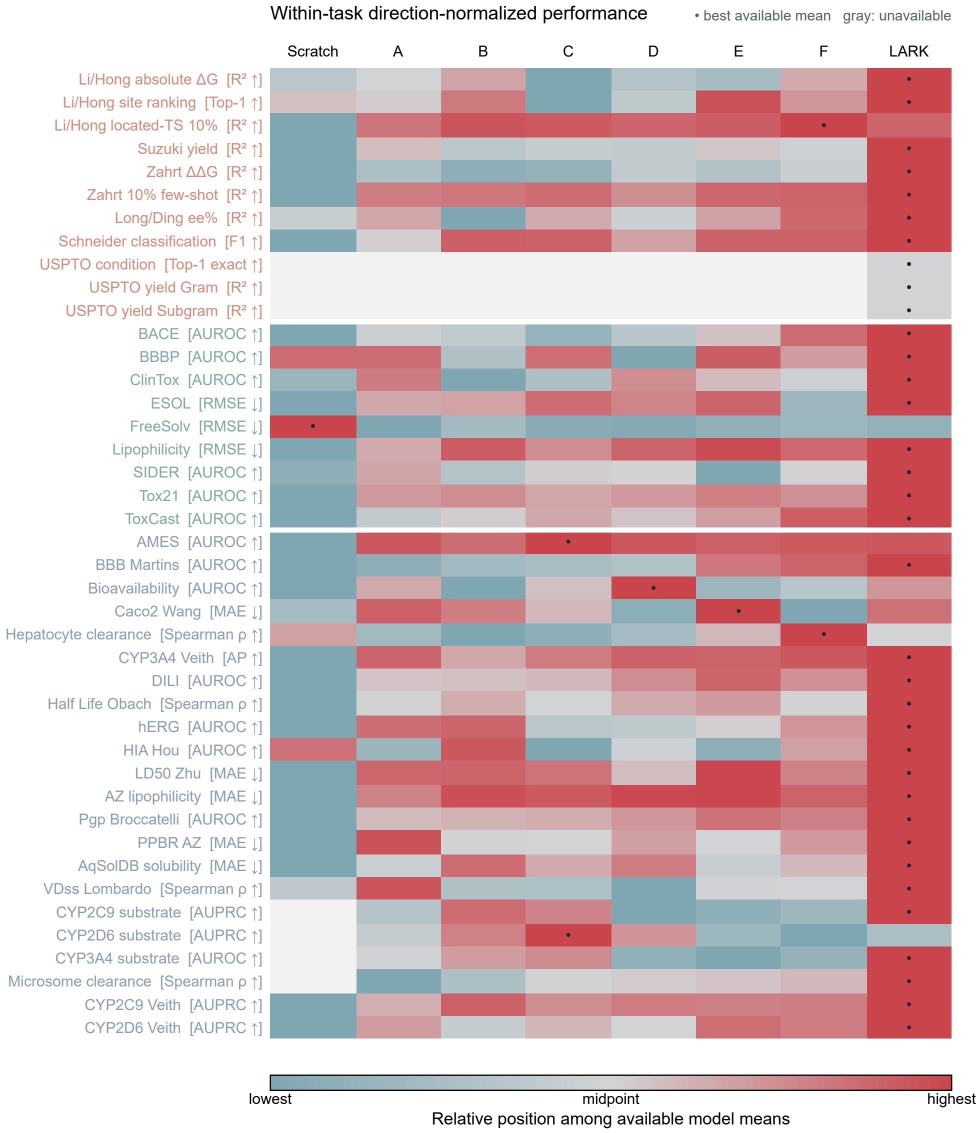

**Figure S2. Full matched-ablation task atlas.** Rows show 42 tasks, ordered as reaction/synthesis tasks, MoleculeNet tasks and TDC ADMET tasks; the colour of each row label identifies its task family. Columns show Scratch, A–F and LARK. Each cell is transformed within its task after reversing lower-is-better metrics: 0 and 1 denote the lowest and highest available model means, respectively. A black dot marks the best available mean, and grey indicates a cell that was not run or was unavailable. Complete model means and sample standard deviations are reported in Supplementary Table S5. A–C are the $\mathrm{R} \to \mathrm{P}$-only variants; D–F and LARK use dual $\mathrm{R} \to \mathrm{P}/\mathrm{P} \to \mathrm{R}$ views.

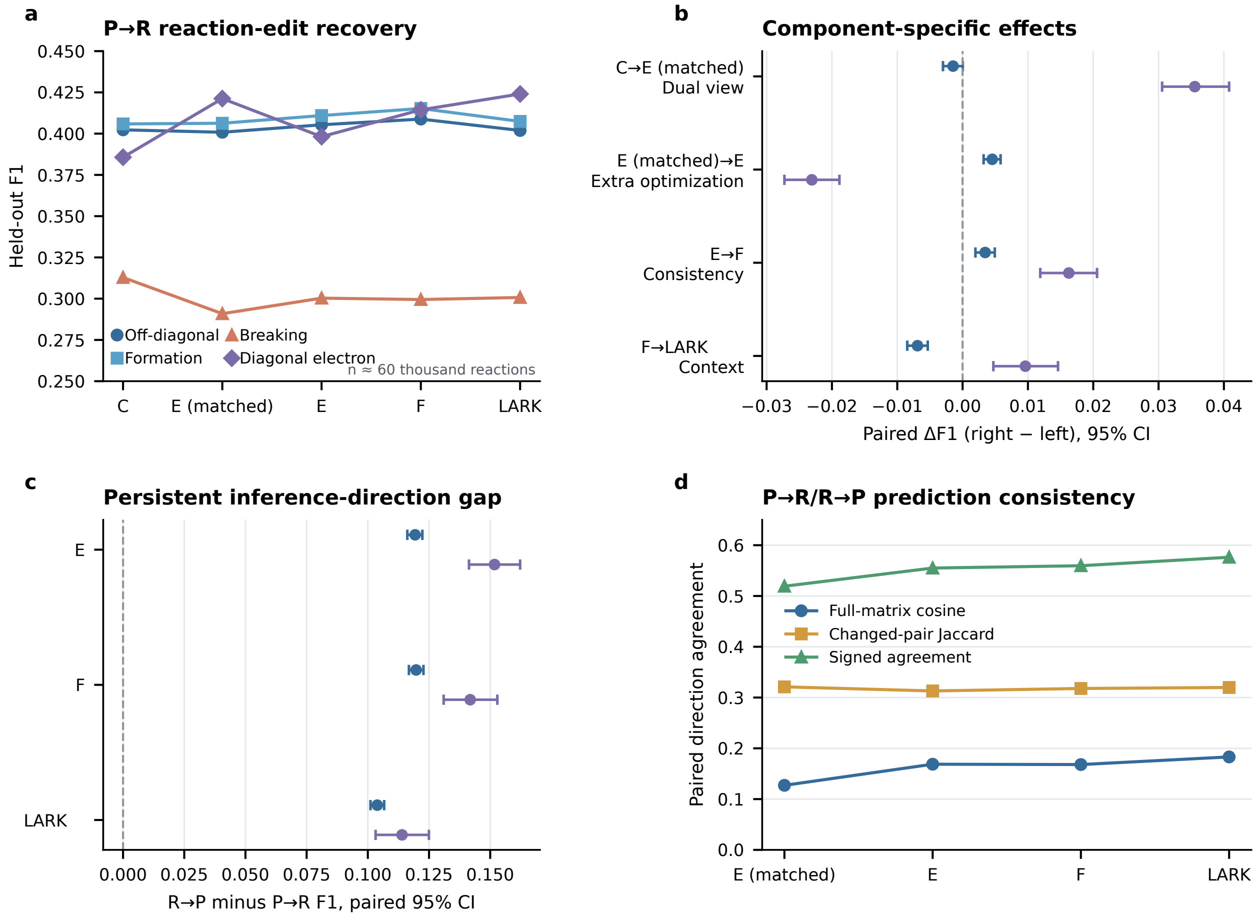

**Figure S3. Directional bond–electron reconstruction and cross-view agreement. a**, Held-out $\Delta\mathrm{BE}$ recovery for off-diagonal, formation, breaking and diagonal electron-change components across the matched pretraining variants in the product-input $\mathrm{P} \to \mathrm{R}$ direction. **b**, Paired changes in component F1 after adding the second view, additional optimization, consistency and reaction context. **c**, Paired difference between $\mathrm{R} \to \mathrm{P}$ and $\mathrm{P} \to \mathrm{R}$ reconstruction. **d**, Full-matrix cosine similarity, changed-pair Jaccard index and signed agreement between the two directional predictions. E(matched) has half the cumulative directional graph-view exposure of E, matching that of C. Points in **b,c** are paired mean differences and error bars are 2,000-resample percentile bootstrap 95% confidence intervals. The analysis includes the complete clean Test collection of approximately 60 thousand reactions.

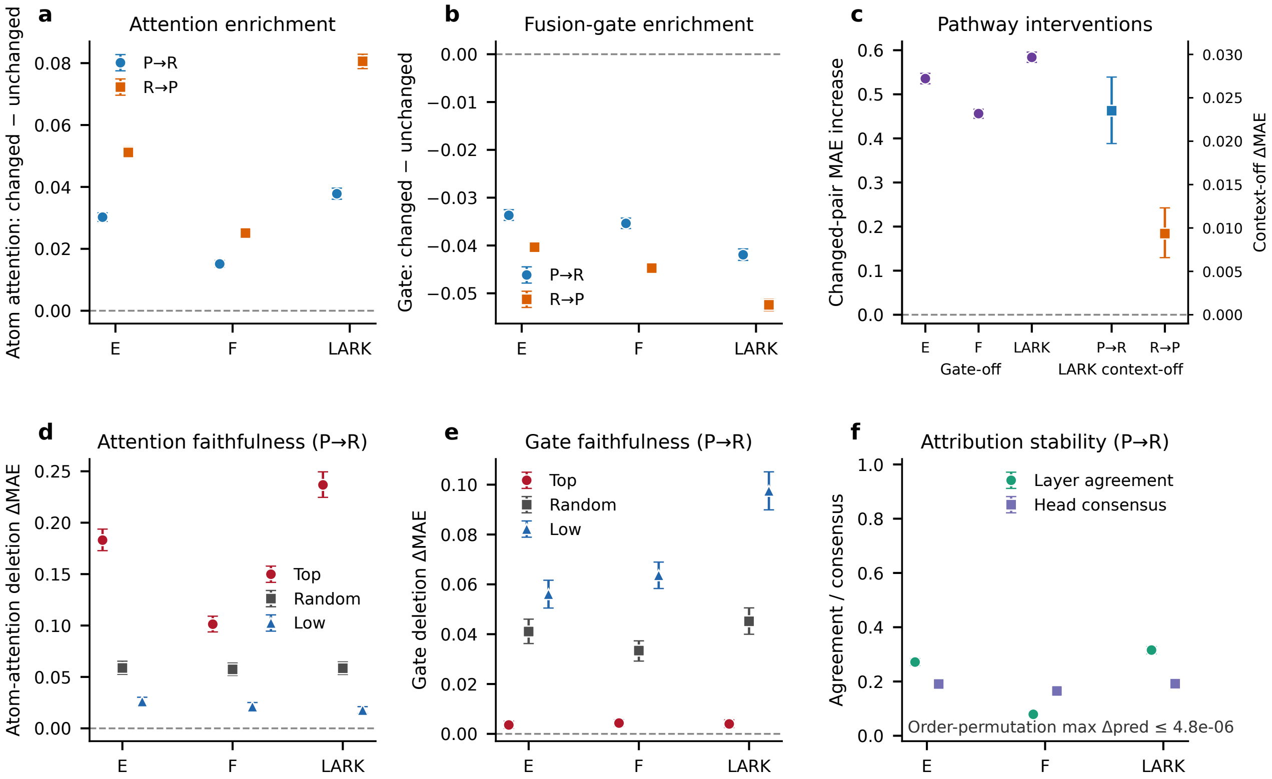

**Figure S4. Attention enrichment, pathway intervention and attribution stability. a,b**, Enrichment of atom attention and fusion-gate magnitude at changed versus unchanged atoms. **c**, Change in changed-pair reconstruction error after suppressing the fusion pathway or context pathway. **d,e**, Change in reconstruction error after suppressing the top-ranked, random or lowest-ranked atom according to atom attention or fusion-gate magnitude. **f**, First-to-last-layer agreement on the top-attended atom and the fraction of final-layer heads selecting the same top atom as the head-averaged attention, together with the 99th percentile of the maximum prediction deviation after atom-order permutation. Points are reaction-level means; error bars are percentile bootstrap 95% confidence intervals.

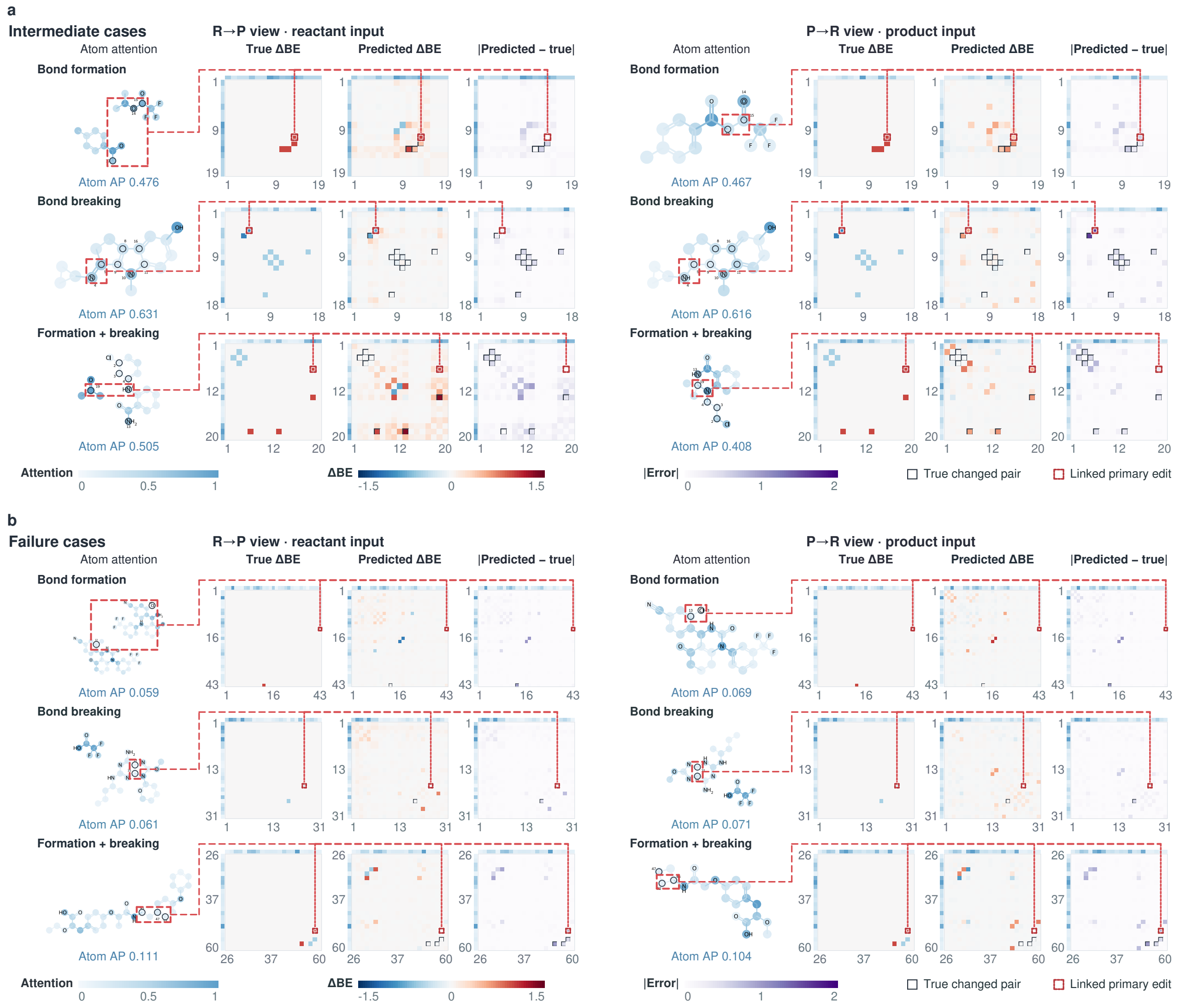

**Figure S5. Intermediate and failure regimes for joint attention and directional $\Delta\mathrm{BE}$ reconstruction. a**, Intermediate cases. **b**, Failure cases. Each panel contains bond formation, bond breaking and mixed formation–breaking reactions evaluated in both the reactant-input $\mathrm{R} \to \mathrm{P}$ and product-input $\mathrm{P} \to \mathrm{R}$ views. Molecular diagrams show atom attention; matrices show the target $\Delta\mathrm{BE}$, predicted $\Delta\mathrm{BE}$ and absolute error. Dashed connectors link the highlighted molecular edit to its atom-pair matrix coordinate. Atom average precision (AP) is annotated beneath each molecular view; the corresponding changed-pair F1 and matrix mean absolute error (MAE) are provided in Source Data. These examples complement the successful cases in Fig. 6d by showing partial recovery and clear failure boundaries.

### E Supplementary Tables

The following five tables are embedded in this document. They contain the reaction-task, MoleculeNet and ADMET baseline comparisons underlying Figs. 2–4, followed by the complete matched-ablation means $\pm$ sample standard deviations used in the supplementary analyses. The matched-ablation means are shown to four decimal places, whereas published comparisons retain their reported precision or are displayed to four decimal places for a common task matrix. Exact plotting values remain in the separately supplied Source Data.

#### Supplementary Table S1. Published baseline comparisons for reaction and synthesis tasks

Reported performance is shown for the ten scalar reaction tasks, followed by the USPTO condition-prediction Top-$k$ comparison. LARK values are taken from the current task matrix and are placed last in each evaluated-task block. Bold type marks the best exact reported mean within each evaluated task. Li/Hong $\Delta\Delta G$ results are provided as related references and are not directly compared with absolute $\Delta G$ prediction. Curve-read values are shown to two decimal places, and the thresholded site-ranking result is shown at its reported threshold; neither is used to assign the bold entry.

| Method | Reported performance |
|---|---|
| **Li/Hong absolute $\Delta G$ ($R^2$ ↑)** | |
| Chemma | 0.9300 |
| PhysOrg-RF (absolute ΔG) | 0.9390 |
| LARK | **0.9828 ± 0.0008** |
| **Related Li/Hong $\Delta\Delta G$ prediction results ($R^2$ ↑)** | |
| SOAP+FP-XGB (ΔΔG) | 0.9680 |
| PhysOrg-RF (ΔΔG) | 0.9630 |
| **Li/Hong site ranking (Top-1 accuracy ↑)** | |
| PhysOrg-RF (direct ΔΔG) | 0.9420 |
| PhysOrg-RF (absolute ΔG → ranking) | 0.8710 |
| Chemma | 0.7874 |
| SOAP+FP-XGB | 0.90 |
| LARK | **0.9598 ± 0.0034** |
| **Li/Hong 10% few-shot ($R^2$ ↑)** | |
| Chemma (10% curve) | 0.52 |
| LARK | **0.8231 ± 0.0602** |
| **Suzuki–Miyaura yield ($R^2$ ↑)** | |
| Chemma | **0.9400** |
| RF + DFT descriptors | 0.9000 |
| DRFP | 0.8500 |
| Yield-BERT | 0.8200 |
| Egret | 0.8500 |
| UGNN | 0.8900 |
| D-MPNN | 0.8500 |
| Rxn Hypergraph | 0.8500 |
| Reaction Graph | 0.8900 |
| T5Chem | 0.8600 |
| RMVP | 0.8500 |
| RMVP (Pretrained) | 0.8600 |
| LARK | 0.8813 ± 0.0182 |
| **Zahrt $\Delta\Delta G$ ($R^2$ ↑)** | |
| Chemma | 0.8900 |
| LARK | **0.8962 ± 0.0190** |
| **Zahrt 10% few-shot ($R^2$ ↑)** | |
| Chemma (10% curve) | 0.74 |
| LARK | **0.7744 ± 0.0226** |
| **Long/Ding ee% ($R^2$ ↑)** | |
| Chemma | **0.9560** |
| FFNN (11×2×1) | 0.9230 |
| FFNN (11×1×1) | 0.9060 |
| Perceptron (11×1) | 0.8450 |
| MLR (11 variables) | 0.7760 |
| SVM (11 variables) | 0.7480 |
| LARK | 0.9335 ± 0.0048 |
| **Schneider reaction classification (Macro-F1 ↑)** | |
| AP3 | 0.9317 ± 0.0031 |
| RxnRep | 0.9533 ± 0.0011 |
| MolR | 0.9389 ± 0.0035 |
| ReaKE | 0.9485 ± 0.0019 |

*Continued on next page*

*Supplementary Table S1 continued*

| Method | Reported performance |
|---|---|
| KAGT | 0.9571 ± 0.0019 |
| LARK | **0.9643 ± 0.0015** |
| **USPTO yield (Gram) ($R^2$ ↑)** | |
| DRFP | **0.1300** |
| Yield-BERT | 0.1170 |
| Egret | 0.1280 |
| UGNN | 0.1170 |
| D-MPNN | 0.1250 |
| Rxn Hypergraph | 0.1180 |
| T5Chem | 0.1160 |
| HRP | 0.1290 |
| LARK | 0.1212 |
| **USPTO yield (Subgram) ($R^2$ ↑)** | |
| DRFP | 0.1970 |
| Yield-BERT | 0.1950 |
| Egret | 0.2060 |
| UGNN | 0.1900 |
| D-MPNN | 0.2020 |
| Rxn Hypergraph | 0.1960 |
| T5Chem | 0.2020 |
| HRP | 0.2000 |
| LARK | **0.2116** |

#### USPTO condition prediction.

| Method | Top-1 ↑ | Top-3 ↑ | Top-5 ↑ | Top-10 ↑ | Top-15 ↑ |
|---|---|---|---|---|---|
| CRM | 0.2600 | 0.3770 | 0.4210 | 0.4610 | 0.4720 |
| Parrot | 0.2690 | 0.4040 | 0.4510 | 0.4910 | 0.5030 |
| AR-GCN | 0.1460 | 0.2370 | 0.2730 | 0.3120 | 0.3260 |
| CIMG | 0.1840 | 0.2710 | 0.3030 | 0.3390 | 0.3530 |
| D-MPNN | 0.1980 | 0.3000 | 0.3340 | 0.3780 | 0.3920 |
| Rxn Hypergraph | 0.2130 | 0.3080 | 0.3450 | 0.3810 | 0.3930 |
| Reaction Graph | **0.3250** | **0.4340** | 0.4720 | 0.5060 | 0.5180 |
| LARK | 0.3015 | 0.4339 | **0.4782** | **0.5167** | **0.5275** |

#### Supplementary Table S2. Task-resolved MoleculeNet classification baseline comparison

Reported test-set performance is shown for LARK and 27 published methods on the MoleculeNet classification tasks. Of these published methods, 25 report all six tasks and are included alongside LARK in the six-task mean comparison in Fig. 4a. Bold type marks the best reported mean for each task. A dash denotes that a value was not reported; mean $\pm$ s.d. is retained where supplied.

| Model | BACE ROC–AUC ↑ | BBBP ROC–AUC ↑ | ClinTox ROC–AUC ↑ | SIDER ROC–AUC ↑ | Tox21 ROC–AUC ↑ | ToxCast ROC–AUC ↑ |
|---|---|---|---|---|---|---|
| ChemBERTa | – | 0.643 | 0.733 | – | – | – |
| MegaMolBART | 0.825 ± 0.008 | 0.689 ± 0.002 | 0.781 ± 0.046 | 0.595 ± 0.018 | 0.739 ± 0.007 | 0.633 ± 0.008 |
| Molformer-XL | 0.821 ± 0.006 | 0.697 ± 0.005 | 0.888 ± 0.032 | 0.625 ± 0.007 | 0.773 ± 0.011 | – |
| EdgePred | 0.646 ± 0.047 | 0.645 ± 0.031 | 0.558 ± 0.062 | 0.567 ± 0.001 | 0.745 ± 0.004 | 0.608 ± 0.005 |
| AttrMask | 0.772 ± 0.014 | 0.702 ± 0.005 | 0.686 ± 0.096 | 0.604 ± 0.006 | 0.742 ± 0.008 | 0.625 ± 0.004 |
| GPT-GNN | 0.776 ± 0.005 | 0.645 ± 0.011 | 0.578 ± 0.042 | 0.575 ± 0.042 | 0.753 ± 0.005 | 0.622 ± 0.001 |
| InfoGraph | 0.739 ± 0.025 | 0.692 ± 0.008 | 0.751 ± 0.050 | 0.592 ± 0.002 | 0.730 ± 0.007 | 0.620 ± 0.003 |
| ContextPred | 0.786 ± 0.014 | 0.712 ± 0.009 | 0.737 ± 0.040 | 0.593 ± 0.014 | 0.733 ± 0.005 | 0.628 ± 0.003 |
| GraphLoG | 0.788 ± 0.007 | 0.678 ± 0.017 | 0.620 ± 0.018 | 0.574 ± 0.023 | 0.730 ± 0.003 | 0.622 ± 0.004 |
| G-Contextual | 0.792 ± 0.003 | 0.703 ± 0.016 | 0.599 ± 0.082 | 0.584 ± 0.006 | 0.752 ± 0.003 | 0.626 ± 0.003 |
| G-Motif | 0.734 ± 0.040 | 0.664 ± 0.034 | 0.778 ± 0.020 | 0.606 ± 0.011 | 0.732 ± 0.008 | 0.626 ± 0.005 |
| GraphCL | 0.687 ± 0.078 | 0.675 ± 0.033 | 0.789 ± 0.042 | 0.601 ± 0.013 | 0.750 ± 0.003 | 0.628 ± 0.002 |
| JOAO | 0.729 ± 0.020 | 0.660 ± 0.006 | 0.663 ± 0.039 | 0.607 ± 0.010 | 0.744 ± 0.007 | 0.627 ± 0.006 |
| MolCLR | 0.711 ± 0.012 | 0.678 ± 0.005 | 0.842 ± 0.015 | 0.587 ± 0.001 | 0.756 ± 0.004 | 0.646 ± 0.001 |
| 3D-InfoMax | 0.794 ± 0.019 | 0.691 ± 0.011 | 0.594 ± 0.032 | 0.534 ± 0.033 | 0.745 ± 0.007 | 0.644 ± 0.009 |
| GraphMVP | 0.812 ± 0.009 | 0.724 ± 0.016 | 0.775 ± 0.042 | 0.639 ± 0.012 | 0.744 ± 0.002 | 0.631 ± 0.004 |
| MOLEBLEND | 0.763 ± 0.003 | 0.665 ± 0.007 | 0.852 ± 0.042 | 0.616 ± 0.015 | 0.769 ± 0.005 | 0.646 ± 0.004 |
| MoleculeSDE | 0.804 ± 0.009 | 0.732 ± 0.005 | 0.866 ± 0.037 | 0.596 ± 0.008 | 0.765 ± 0.003 | 0.652 ± 0.003 |
| KV-PLM | 0.785 ± 0.027 | 0.705 ± 0.005 | 0.892 ± 0.027 | 0.598 ± 0.006 | 0.721 ± 0.010 | 0.550 ± 0.017 |
| MoleculeSTM1 | 0.820 ± 0.004 | 0.708 ± 0.019 | 0.866 ± 0.023 | 0.637 ± 0.008 | 0.757 ± 0.009 | 0.652 ± 0.004 |
| MoleculeSTM2 | 0.808 ± 0.013 | 0.700 ± 0.005 | 0.925 ± 0.011 | 0.610 ± 0.011 | 0.769 ± 0.005 | 0.651 ± 0.004 |
| MoMu | 0.767 ± 0.021 | 0.705 ± 0.020 | 0.799 ± 0.041 | 0.605 ± 0.009 | 0.756 ± 0.003 | 0.634 ± 0.005 |
| MolFM | 0.839 ± 0.011 | 0.729 ± 0.001 | 0.797 ± 0.016 | 0.642 ± 0.009 | 0.772 ± 0.007 | 0.644 ± 0.002 |
| MEMO | 0.826 ± 0.003 | 0.716 ± 0.010 | 0.816 ± 0.037 | 0.612 ± 0.006 | 0.767 ± 0.004 | 0.649 ± 0.008 |
| M2UMol | 0.843 ± 0.014 | 0.682 ± 0.006 | 0.922 ± 0.005 | 0.660 ± 0.009 | 0.762 ± 0.001 | 0.663 ± 0.005 |
| ChemRL-GEM | 0.813 ± 0.011 | 0.679 ± 0.013 | 0.882 ± 0.009 | 0.603 ± 0.008 | 0.767 ± 0.006 | 0.673 ± 0.004 |
| Uni-Mol | 0.776 ± 0.059 | 0.700 ± 0.007 | 0.862 ± 0.016 | 0.629 ± 0.011 | 0.785 ± 0.031 | **0.689 ± 0.002** |
| LARK | **0.8530 ± 0.0188** | **0.7339 ± 0.0075** | **0.9431 ± 0.0094** | **0.6655 ± 0.0037** | **0.7940 ± 0.0034** | 0.6844 ± 0.0022 |

#### Supplementary Table S3. Task-resolved MoleculeNet regression baseline comparison

Reported test-set performance is shown for the MoleculeNet regression tasks. Bold type marks the best reported mean for each task. A dash denotes that a value was not reported; mean ± s.d. is retained where supplied.

| Model | ESOL<br>RMSE ↓ | FreeSolv<br>RMSE ↓ | Lipophilicity<br>RMSE ↓ |
|---|---|---|---|
| AttrMask | 1.112 | – | 0.730 |
| ContextPred | 1.196 | – | 0.702 |
| JOAO | 1.120 | – | 0.708 |
| 3D-InfoMax | 0.894 | 2.337 | 0.695 |
| GraphMVP | 1.029 | – | 0.681 |
| MEMO | 0.984 | – | 0.707 |
| M2UMol | 0.914 | **1.894** | 0.667 |
| LARK | **0.8089 ± 0.0301** | 2.5366 ± 0.1018 | **0.6397 ± 0.0073** |

#### Supplementary Table S4. Absolute performance across 22 TDC ADMET tasks

Values are reported means ± s.d. for MolE, the fixed all-atom-QM QM-Graphormer configuration and LARK. Bold type marks the best reported mean for each task after accounting for metric direction. These absolute values underlie the direction-normalized pairwise margins in Fig. 4b.

| Task | Metric | MolE | QM-Graphormer | LARK |
|---|---|---|---|---|
| **Absorption** | | | | |
| Caco-2 | MAE ↓ | 0.3290 ± 0.0080 | 0.3540 ± 0.0150 | **0.3253 ± 0.0294** |
| HIA | AUROC ↑ | 0.9840 ± 0.0050 | 0.9820 ± 0.0030 | **0.9847 ± 0.0037** |
| P-gp | AUROC ↑ | 0.9300 ± 0.0050 | 0.9130 ± 0.0150 | **0.9340 ± 0.0087** |
| Bioavailability | AUROC ↑ | 0.6400 ± 0.0460 | 0.6730 ± 0.0280 | **0.6907 ± 0.0512** |
| Lipophilicity | MAE ↓ | 0.4060 ± 0.0090 | **0.3930 ± 0.0050** | 0.4268 ± 0.0074 |
| Solubility | MAE ↓ | 0.7760 ± 0.0190 | **0.7200 ± 0.0100** | 0.7328 ± 0.0074 |
| **Distribution** | | | | |
| BBB | AUROC ↑ | **0.9030 ± 0.0030** | 0.8720 ± 0.0210 | 0.9013 ± 0.0137 |
| PPBR | MAE ↓ | **7.2290 ± 0.1680** | 7.5890 ± 0.2030 | 7.7375 ± 0.3172 |
| VDss | Spearman ↑ | **0.6440 ± 0.0130** | 0.6240 ± 0.0200 | 0.5620 ± 0.0417 |
| **Metabolism** | | | | |
| CYP2D6 inhibition | AUPRC ↑ | 0.6790 ± 0.0060 | 0.6210 ± 0.0460 | **0.6932 ± 0.0095** |
| CYP3A4 inhibition | AUPRC ↑ | 0.8760 ± 0.0020 | 0.7970 ± 0.0290 | **0.8818 ± 0.0037** |
| CYP2C9 inhibition | AUPRC ↑ | 0.7820 ± 0.0010 | 0.7030 ± 0.0220 | **0.7912 ± 0.0093** |
| CYP2D6 substrate | AUPRC ↑ | **0.6920 ± 0.0170** | 0.6480 ± 0.0310 | 0.6762 ± 0.0402 |
| CYP3A4 substrate | AUROC ↑ | **0.6920 ± 0.0190** | 0.6300 ± 0.0150 | 0.6245 ± 0.0317 |
| CYP2C9 substrate | AUPRC ↑ | 0.4090 ± 0.0140 | 0.3740 ± 0.0280 | **0.4237 ± 0.0313** |
| **Excretion** | | | | |
| Half-life | Spearman ↑ | **0.5780 ± 0.0320** | 0.4620 ± 0.1540 | 0.4108 ± 0.0469 |
| Microsomal clearance | Spearman ↑ | **0.6320 ± 0.0080** | 0.5480 ± 0.0290 | 0.5914 ± 0.0201 |
| Hepatocyte clearance | Spearman ↑ | **0.4560 ± 0.0270** | 0.3820 ± 0.0320 | 0.4120 ± 0.0255 |
| **Toxicity** | | | | |
| hERG | AUROC ↑ | 0.8350 ± 0.0180 | 0.7880 ± 0.0290 | **0.8356 ± 0.0206** |
| AMES | AUROC ↑ | 0.8340 ± 0.0150 | 0.8220 ± 0.0050 | **0.8479 ± 0.0060** |
| DILI | AUROC ↑ | 0.8520 ± 0.0220 | 0.8920 ± 0.0330 | **0.9237 ± 0.0229** |
| LD50 | MAE ↓ | 0.6020 ± 0.0160 | **0.5590 ± 0.0160** | 0.5746 ± 0.0205 |

#### Supplementary Table S5. Complete matched-ablation means and s.d. across 42 tasks

Values are means $\pm$ sample standard deviations across downstream repeats. Reaction and MoleculeNet tasks use three downstream repeats unless noted; USPTO condition prediction has $n = 1$, and the Subgram repeat count is unavailable. TDC ADMET tasks use five frozen split identities. The LARK entries for USPTO condition prediction and Gram- and Subgram-scale yield are shown as means only because no standard deviation is available in the task matrix. Bold type marks the best available mean within each task after accounting for metric direction. A dash denotes an unavailable or unrun cell.

##### Reaction and synthesis tasks.

| Task | Metric | Scratch | A | B | C | D | E | F | LARK |
|---|---|---|---|---|---|---|---|---|---|
| Li/Hong absolute $\Delta G$ | $R^2 \uparrow$ | 0.9754 $\pm$ 0.0034 | 0.9770 $\pm$ 0.0075 | 0.9790 $\pm$ 0.0014 | 0.9713 $\pm$ 0.0012 | 0.9749 $\pm$ 0.0042 | 0.9740 $\pm$ 0.0028 | 0.9787 $\pm$ 0.0022 | **0.9828 $\pm$ 0.0008** |
| Li/Hong site ranking | Top-1 accuracy $\uparrow$ | 0.9510 $\pm$ 0.0017 | 0.9500 $\pm$ 0.0059 | 0.9559 $\pm$ 0.0106 | 0.9392 $\pm$ 0.0045 | 0.9471 $\pm$ 0.0078 | 0.9588 $\pm$ 0.0051 | 0.9539 $\pm$ 0.0111 | **0.9598 $\pm$ 0.0034** |
| Li/Hong 10% few-shot | $R^2 \uparrow$ | 0.5753 $\pm$ 0.3162 | 0.8063 $\pm$ 0.0217 | 0.8379 $\pm$ 0.0424 | 0.8332 $\pm$ 0.0530 | 0.8223 $\pm$ 0.0366 | 0.8293 $\pm$ 0.0262 | **0.8550 $\pm$ 0.0055** | 0.8231 $\pm$ 0.0602 |
| Suzuki–Miyaura yield | $R^2 \uparrow$ | 0.8440 $\pm$ 0.0135 | 0.8656 $\pm$ 0.0125 | 0.8573 $\pm$ 0.0144 | 0.8593 $\pm$ 0.0020 | 0.8588 $\pm$ 0.0253 | 0.8644 $\pm$ 0.0206 | 0.8612 $\pm$ 0.0220 | **0.8813 $\pm$ 0.0182** |
| Zahrt $\Delta\Delta G$ | $R^2 \uparrow$ | 0.8487 $\pm$ 0.0237 | 0.8613 $\pm$ 0.0217 | 0.8530 $\pm$ 0.0116 | 0.8544 $\pm$ 0.0431 | 0.8675 $\pm$ 0.0079 | 0.8628 $\pm$ 0.0098 | 0.8696 $\pm$ 0.0073 | **0.8962 $\pm$ 0.0190** |
| Zahrt 10% few-shot | $R^2 \uparrow$ | 0.1588 $\pm$ 0.3259 | 0.6496 $\pm$ 0.1161 | 0.6629 $\pm$ 0.0730 | 0.6901 $\pm$ 0.0569 | 0.6112 $\pm$ 0.2324 | 0.6997 $\pm$ 0.0428 | 0.7053 $\pm$ 0.0114 | **0.7744 $\pm$ 0.0226** |
| Long/Ding ee% | $R^2 \uparrow$ | 0.9191 $\pm$ 0.0169 | 0.9247 $\pm$ 0.0129 | 0.9078 $\pm$ 0.0325 | 0.9244 $\pm$ 0.0092 | 0.9193 $\pm$ 0.0185 | 0.9252 $\pm$ 0.0129 | 0.9304 $\pm$ 0.0060 | **0.9335 $\pm$ 0.0048** |
| Schneider reaction classification | Macro-F1 $\uparrow$ | 0.9567 $\pm$ 0.0014 | 0.9607 $\pm$ 0.0033 | 0.9636 $\pm$ 0.0020 | 0.9636 $\pm$ 0.0003 | 0.9618 $\pm$ 0.0028 | 0.9635 $\pm$ 0.0024 | 0.9635 $\pm$ 0.0041 | **0.9643 $\pm$ 0.0015** |
| USPTO condition prediction | Beam Top-1 exact accuracy $\uparrow$ | – | – | – | – | – | – | – | **0.3015** |
| USPTO yield (Gram) | $R^2 \uparrow$ | – | – | – | – | – | – | – | **0.1212** |
| USPTO yield (Subgram) | $R^2 \uparrow$ | – | – | – | – | – | – | – | **0.2116** |

##### MoleculeNet tasks.

| Task | Metric | Scratch | A | B | C | D | E | F | LARK |
|---|---|---|---|---|---|---|---|---|---|
| BACE | ROC–AUC $\uparrow$ | 0.8203 $\pm$ 0.0301 | 0.8351 $\pm$ 0.0199 | 0.8331 $\pm$ 0.0307 | 0.8250 $\pm$ 0.0173 | 0.8317 $\pm$ 0.0212 | 0.8388 $\pm$ 0.0014 | 0.8485 $\pm$ 0.0108 | **0.8530 $\pm$ 0.0188** |
| BBBP | ROC–AUC $\uparrow$ | 0.7308 $\pm$ 0.0046 | 0.7306 $\pm$ 0.0112 | 0.7178 $\pm$ 0.0208 | 0.7305 $\pm$ 0.0136 | 0.7113 $\pm$ 0.0394 | 0.7318 $\pm$ 0.0049 | 0.7269 $\pm$ 0.0073 | **0.7339 $\pm$ 0.0075** |
| ClinTox | ROC–AUC $\uparrow$ | 0.8963 $\pm$ 0.0239 | 0.9324 $\pm$ 0.0208 | 0.8876 $\pm$ 0.0129 | 0.9029 $\pm$ 0.0208 | 0.9289 $\pm$ 0.0053 | 0.9203 $\pm$ 0.0192 | 0.9130 $\pm$ 0.0205 | **0.9431 $\pm$ 0.0094** |
| ESOL | RMSE $\downarrow$ | 0.8923 $\pm$ 0.0160 | 0.8378 $\pm$ 0.0128 | 0.8367 $\pm$ 0.0049 | 0.8208 $\pm$ 0.0150 | 0.8280 $\pm$ 0.0161 | 0.8182 $\pm$ 0.0118 | 0.8765 $\pm$ 0.0299 | **0.8089 $\pm$ 0.0301** |
| FreeSolv | RMSE $\downarrow$ | **1.9908 $\pm$ 0.0419** | 2.6102 $\pm$ 0.0857 | 2.4761 $\pm$ 0.1342 | 2.5744 $\pm$ 0.1219 | 2.5927 $\pm$ 0.0286 | 2.5403 $\pm$ 0.2200 | 2.4996 $\pm$ 0.1224 | 2.5366 $\pm$ 0.1018 |
| Lipophilicity | RMSE $\downarrow$ | 0.7464 $\pm$ 0.0034 | 0.6779 $\pm$ 0.0037 | 0.6491 $\pm$ 0.0151 | 0.6668 $\pm$ 0.0259 | 0.6518 $\pm$ 0.0174 | 0.6430 $\pm$ 0.0085 | 0.6538 $\pm$ 0.0054 | **0.6397 $\pm$ 0.0073** |
| SIDER | ROC–AUC $\uparrow$ | 0.6181 $\pm$ 0.0143 | 0.6478 $\pm$ 0.0107 | 0.6297 $\pm$ 0.0288 | 0.6406 $\pm$ 0.0138 | 0.6396 $\pm$ 0.0027 | 0.6131 $\pm$ 0.0157 | 0.6400 $\pm$ 0.0021 | **0.6655 $\pm$ 0.0037** |
| Tox21 | ROC–AUC $\uparrow$ | 0.7393 $\pm$ 0.0044 | 0.7775 $\pm$ 0.0040 | 0.7799 $\pm$ 0.0060 | 0.7753 $\pm$ 0.0018 | 0.7780 $\pm$ 0.0045 | 0.7826 $\pm$ 0.0030 | 0.7794 $\pm$ 0.0028 | **0.7940 $\pm$ 0.0034** |
| ToxCast | ROC–AUC $\uparrow$ | 0.6561 $\pm$ 0.0131 | 0.6671 $\pm$ 0.0044 | 0.6709 $\pm$ 0.0048 | 0.6746 $\pm$ 0.0034 | 0.6719 $\pm$ 0.0015 | 0.6754 $\pm$ 0.0093 | 0.6817 $\pm$ 0.0017 | **0.6844 $\pm$ 0.0022** |

##### TDC ADMET tasks.

| Task | Metric | Scratch | A | B | C | D | E | F | LARK |
|---|---|---|---|---|---|---|---|---|---|
| AMES | ROC–AUC $\uparrow$ | 0.8035 $\pm$ 0.0093 | 0.8482 $\pm$ 0.0106 | 0.8443 $\pm$ 0.0087 | **0.8514 $\pm$ 0.0090** | 0.8479 $\pm$ 0.0105 | 0.8467 $\pm$ 0.0102 | 0.8476 $\pm$ 0.0077 | 0.8479 $\pm$ 0.0060 |
| BBB Martins | ROC–AUC $\uparrow$ | 0.8806 $\pm$ 0.0236 | 0.8824 $\pm$ 0.0127 | 0.8848 $\pm$ 0.0131 | 0.8858 $\pm$ 0.0146 | 0.8869 $\pm$ 0.0168 | 0.8976 $\pm$ 0.0185 | 0.8990 $\pm$ 0.0124 | **0.9013 $\pm$ 0.0137** |
| Bioavailability Ma | ROC–AUC $\uparrow$ | 0.6406 $\pm$ 0.0190 | 0.6860 $\pm$ 0.0252 | 0.6402 $\pm$ 0.0314 | 0.6804 $\pm$ 0.0465 | **0.7112 $\pm$ 0.0205** | 0.6527 $\pm$ 0.0529 | 0.6641 $\pm$ 0.0144 | 0.6907 $\pm$ 0.0512 |
| Caco2 Wang | MAE $\downarrow$ | 0.3730 $\pm$ 0.0315 | 0.3206 $\pm$ 0.0257 | 0.3291 $\pm$ 0.0397 | 0.3450 $\pm$ 0.0506 | 0.3860 $\pm$ 0.0392 | **0.3127 $\pm$ 0.0146** | 0.3925 $\pm$ 0.0447 | 0.3253 $\pm$ 0.0294 |
| Clearance Hepatocyte AZ | Spearman's $\rho \uparrow$ | 0.4222 $\pm$ 0.0628 | 0.3966 $\pm$ 0.0643 | 0.3856 $\pm$ 0.0205 | 0.3910 $\pm$ 0.0173 | 0.3987 $\pm$ 0.0421 | 0.4180 $\pm$ 0.0317 | **0.4395 $\pm$ 0.0213** | 0.4120 $\pm$ 0.0255 |
| CYP3A4 Veith | Average precision $\uparrow$ | 0.8463 $\pm$ 0.0106 | 0.8776 $\pm$ 0.0040 | 0.8695 $\pm$ 0.0055 | 0.8748 $\pm$ 0.0039 | 0.8780 $\pm$ 0.0046 | 0.8779 $\pm$ 0.0016 | 0.8792 $\pm$ 0.0069 | **0.8818 $\pm$ 0.0037** |
| DILI | ROC–AUC $\uparrow$ | 0.8741 $\pm$ 0.0166 | 0.9015 $\pm$ 0.0233 | 0.9023 $\pm$ 0.0286 | 0.9036 $\pm$ 0.0277 | 0.9110 $\pm$ 0.0300 | 0.9181 $\pm$ 0.0138 | 0.9105 $\pm$ 0.0369 | **0.9237 $\pm$ 0.0229** |
| Half Life Obach | Spearman's $\rho \uparrow$ | 0.2634 $\pm$ 0.0261 | 0.3385 $\pm$ 0.0895 | 0.3568 $\pm$ 0.0303 | 0.3363 $\pm$ 0.0824 | 0.3586 $\pm$ 0.0952 | 0.3661 $\pm$ 0.0659 | 0.3381 $\pm$ 0.0342 | **0.4108 $\pm$ 0.0469** |
| hERG | ROC–AUC $\uparrow$ | 0.7962 $\pm$ 0.0664 | 0.8298 $\pm$ 0.0180 | 0.8311 $\pm$ 0.0338 | 0.8102 $\pm$ 0.0474 | 0.8112 $\pm$ 0.0288 | 0.8167 $\pm$ 0.0237 | 0.8245 $\pm$ 0.0339 | **0.8356 $\pm$ 0.0206** |

*Continued on next page*

*Supplementary Table S5, TDC ADMET tasks continued*

| Task | Metric | Scratch | A | B | C | D | E | F | LARK |
|---|---|---|---|---|---|---|---|---|---|
| HIA Hou | ROC–AUC ↑ | 0.9834 ± 0.0086 | 0.9780 ± 0.0021 | 0.9841 ± 0.0068 | 0.9767 ± 0.0070 | 0.9805 ± 0.0065 | 0.9775 ± 0.0182 | 0.9821 ± 0.0050 | **0.9847 ± 0.0037** |
| LD50 Zhu | MAE ↓ | 0.6532 ± 0.0193 | 0.5841 ± 0.0296 | 0.5837 ± 0.0326 | 0.5880 ± 0.0172 | 0.6080 ± 0.0171 | 0.5754 ± 0.0130 | 0.5917 ± 0.0165 | **0.5746 ± 0.0205** |
| Lipophilicity AstraZeneca | MAE ↓ | 0.5293 ± 0.0270 | 0.4489 ± 0.0059 | 0.4309 ± 0.0120 | 0.4345 ± 0.0093 | 0.4286 ± 0.0054 | 0.4281 ± 0.0094 | 0.4383 ± 0.0059 | **0.4268 ± 0.0074** |
| Pgp Broccatelli | ROC–AUC ↑ | 0.8976 ± 0.0101 | 0.9189 ± 0.0129 | 0.9202 ± 0.0120 | 0.9207 ± 0.0081 | 0.9234 ± 0.0121 | 0.9279 ± 0.0054 | 0.9263 ± 0.0090 | **0.9340 ± 0.0087** |
| PPBR AZ | MAE ↓ | 8.0936 ± 0.1907 | 7.7565 ± 0.1823 | 7.9138 ± 0.2348 | 7.9186 ± 0.3029 | 7.8510 ± 0.2581 | 7.9158 ± 0.2360 | 7.8444 ± 0.3511 | **7.7375 ± 0.3172** |
| Solubility AqSolDB | MAE ↓ | 0.7823 ± 0.0058 | 0.7600 ± 0.0073 | 0.7398 ± 0.0173 | 0.7504 ± 0.0194 | 0.7428 ± 0.0207 | 0.7610 ± 0.0227 | 0.7529 ± 0.0113 | **0.7328 ± 0.0074** |
| VDss Lombardo | Spearman's $\rho$ ↑ | 0.5381 ± 0.0273 | 0.5597 ± 0.0520 | 0.5348 ± 0.0645 | 0.5344 ± 0.0491 | 0.5237 ± 0.0533 | 0.5423 ± 0.0606 | 0.5429 ± 0.0431 | **0.5620 ± 0.0417** |
| CYP2C9 Substrate CarbonMangels | AUPRC ↑ | – | 0.4002 ± 0.0441 | 0.4187 ± 0.0473 | 0.4160 ± 0.0450 | 0.3888 ± 0.0158 | 0.3923 ± 0.0149 | 0.3951 ± 0.0301 | **0.4237 ± 0.0313** |
| CYP2D6 Substrate CarbonMangels | AUPRC ↑ | – | 0.6851 ± 0.0078 | 0.7065 ± 0.0190 | **0.7189 ± 0.0200** | 0.7027 ± 0.0134 | 0.6717 ± 0.0306 | 0.6610 ± 0.0282 | 0.6762 ± 0.0402 |
| CYP3A4 Substrate CarbonMangels | AUROC ↑ | – | 0.6087 ± 0.0209 | 0.6152 ± 0.0258 | 0.6169 ± 0.0077 | 0.5972 ± 0.0197 | 0.5940 ± 0.0120 | 0.5980 ± 0.0086 | **0.6245 ± 0.0317** |
| Clearance Microsome AZ | Spearman's $\rho$ ↑ | – | 0.5404 ± 0.0290 | 0.5540 ± 0.0383 | 0.5661 ± 0.0210 | 0.5673 ± 0.0157 | 0.5686 ± 0.0237 | 0.5709 ± 0.0355 | **0.5914 ± 0.0201** |
| CYP2C9 Veith | AUPRC ↑ | 0.7418 ± 0.0158 | 0.7730 ± 0.0060 | 0.7863 ± 0.0107 | 0.7785 ± 0.0093 | 0.7815 ± 0.0073 | 0.7807 ± 0.0113 | 0.7812 ± 0.0099 | **0.7912 ± 0.0093** |
| CYP2D6 Veith | AUPRC ↑ | 0.6551 ± 0.0201 | 0.6814 ± 0.0097 | 0.6711 ± 0.0145 | 0.6781 ± 0.0175 | 0.6740 ± 0.0159 | 0.6874 ± 0.0140 | 0.6855 ± 0.0144 | **0.6932 ± 0.0095** |
